# The E525K β-Myosin Mutation Causes Hypocontractility in Cardiomyocytes Without Altering Crossbridge Cycling

**DOI:** 10.64898/2026.06.18.733270

**Authors:** Kalen Z. Robeson, Timothy S. McMillen, Kristina Kooiker, Kerry Y. Kao, Travis C. Tune, Kieran Fruebis, Michael A. Geeves, Andrew P. Wescott, Rachelle Soriano, Alex J. Goldstein, Matthew C. Childers, Rama Reddy Goluguri, Divya Pathak, Åshild Telle, Iselin Crosby, Nathan J. Sniadecki, Joseph D. Powers, Jennifer Davis, Farid Moussavi-Harami, James A. Spudich, Kathleen M. Ruppel, Michael Regnier

## Abstract

The cardiac β-myosin (MYH7) variant E525K was identified in 2012 in a patient with dilated cardiomyopathy (DCM). Work using engineered myosin constructs has shown that this variant stabilizes the interacting heads motif (IHM) of myosin and increases the ATPase activity of variant motor S1 heads. However, no measurements have been made in myofilaments or cardiomyocytes to determine its effect on contractile function. Here, we present force and contractile kinetics measurements from induced pluripotent stem cell-derived cardiomyocytes (iPSC-CMs) gene-edited for heterozygous expression of the E525K variant. Single-cell contraction for E525K hiPSC-CMs was reduced by 65%, and isometric twitch force in engineered heart tissues (EHTs) was reduced by 39%. In contrast, maximal Ca^2+^ activated isometric force in isolated myofibrils was increased by 45%, and sub-maximal Ca^2+^ activated force was similar to that of WT myofibrils. Structural analysis revealed reduced myofibril content (13.7% decrease) and decreased organization (increased z-disk dispersion angle) in E525K cells. We confirmed that E525K S1 myosin has higher actin affinity than WT S1 and elevated ATPase activity. However, there was no change in S1 ADP release rate. There was also no change in either the rate of force development or relaxation in myofibrils, cells, or EHTs. These findings suggest that E525K myosin crossbridge cycling is not altered during loaded contractions. To understand how the twitch force of myocytes decreased while the maximal isometric force increased, we used a spatially explicit, stochastic-kinetic sarcomere model. The results were explained by changes in three rates: reduced recruitment from the OFF/IHM state and increased rates of actin binding and Pi release. Finite element modeling was used to demonstrate that additional contractile deficits in cells and EHTs likely result from myofibrillar disorganization. This study demonstrates the value of multi-scale analysis and coupled computational modeling for understanding the molecular mechanisms underlying sarcomere mutations in cardiomyocytes.

**Graphical Abstract:** Graphical Abstract:
A model for how the hypothesized effects of the E525K mutation impact contracting myofibrils Here, we show that the E525K mutation impacts contraction in multiple ways: (1) Decreased sarcomere organization in cells and tissues drives decreased force generation. (2) Increased binding affinity of E525K myosin for actin and faster Pi release contribute to increased force generation in isolated myofibrils. (3) Increased IHM stability reduces twitch force. (4) The rate-limiting step of loaded crossbridge cycling, ADP release, is unchanged by the E525K mutation, and the rates of loaded contraction and relaxation are unchanged at all scales of contraction. Finally, a spatially explicit sarcomere model is used to demonstrate that these biochemical changes to myosin recruitment, binding, and cycling can explain force generation at tissue, cell, and myofibril levels.

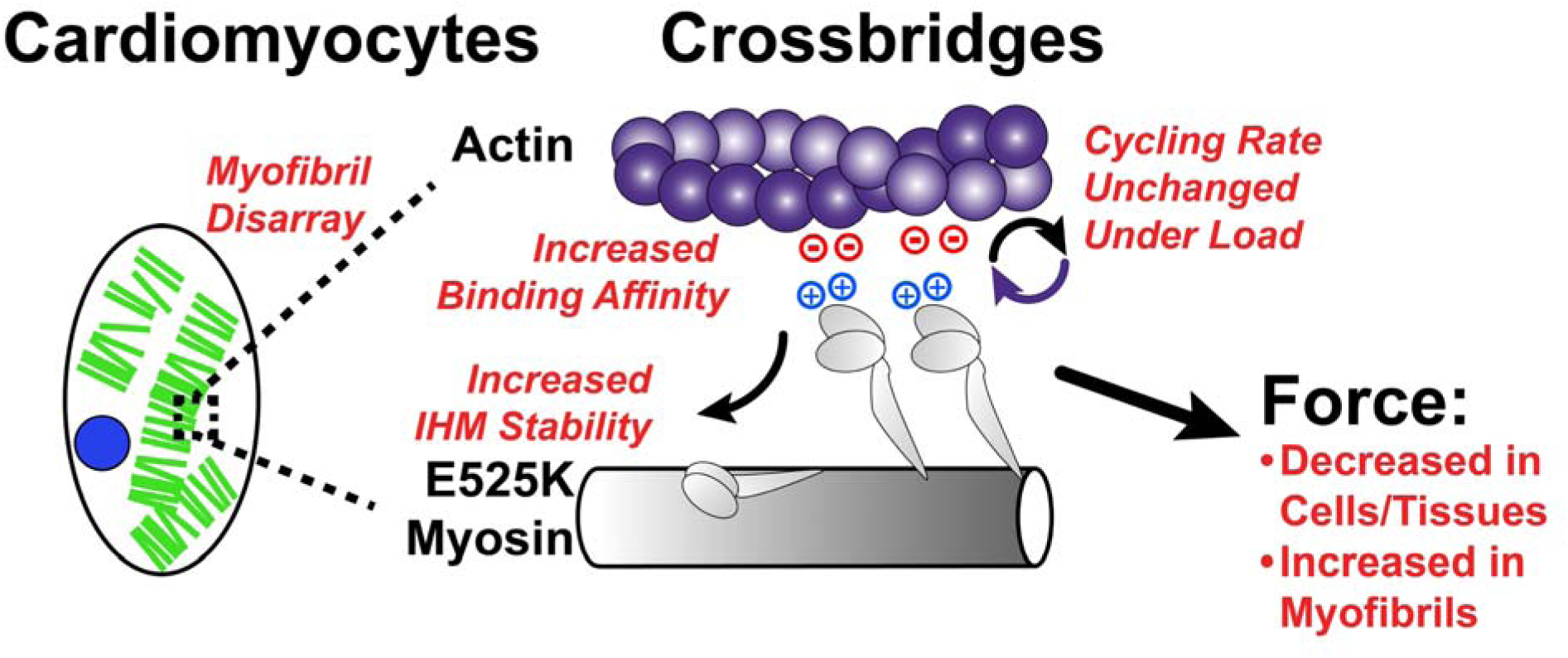

## Introduction

Mutations in sarcomere proteins can be causal for the initiation and progression of familial forms of hypertrophic cardiomyopathy (HCM) and dilated cardiomyopathy (DCM)[1]. This has led to increased genetic screening of cardiomyopathy patients, but the predictive power of these screens is limited by our understanding of how individual mutations alter cardiac function. Identifying the tension-time integral as a unifying determinant of whether a mutation drives HCM or DCM has improved this forecasting[2,3], as have biophysical and structural studies of identified hot spot regions in sarcomere proteins such as myosin[4–6]. However, our understanding of mutation-specific disease mechanisms is far from complete, and continued efforts are needed to elucidate how molecular changes in protein structure and function scale up to affect organized muscle structure at the sarcomere, cell, and tissue levels.

The heterozygous E525K mutation in MYH7 was first identified through a genetic screen of patients with DCM in a single person diagnosed with DCM. However, the mechanisms by which this mutation might elicit and contribute to the DCM phenotype remained unclear[7]. More recently, a genetic screen of patients with HCM identified a family carrying the same mutation[8]. Importantly, additional sarcomere mutations were found in this individual, further complicating interpretations of how E525K may alter myosin behavior. Recent structural and biochemical studies have used isolated myosin constructs to study the E525K mutation[9–12]. These reports indicate that the E525K mutation increases myosin motor domain (Sub-fragment 1, S1) ATPase activity but decreases the ATPase activity of 2-headed myosin constructs (Heavy Meromyosin, HMM) in both the presence and absence of actin[9,10]. The increased S1 ATPase activity was attributed to an increase in actin affinity and phosphate release rate for E525K myosin[11], while the decreased HMM ATPase activity was attributed to stabilization of the myosin OFF state[9,10]. While these biochemical studies have been insightful, the E525K myosin variant has not been studied in intact myofilaments during loaded contraction, where it may influence recruitment from the thick filament backbone, binding to actin, and/or actin-myosin crossbridge cycling (**Figure 1A and B**). Under conditions of high load, one or more of these myosin functions may be affected by the mutation. For example, during isometric contraction, the release of ADP from the myosins is significantly slowed[13,14], switching the rate-limiting step in the acto-myosin chemomechanical cycle from phosphate release to ADP release (**Figure 1B**). Additionally, the recruitment of myosin heads from the thick filament backbone in sarcomeres is facilitated by increased thick filament strain[15]. Since recruitment and phosphate release are two key mechanisms by which the E525K mutation has been hypothesized to affect myosin activity[9–12] (**Figure 1B**), it is important to confirm whether this is consistent in intact myofilaments during contraction.

**Figure 1:**
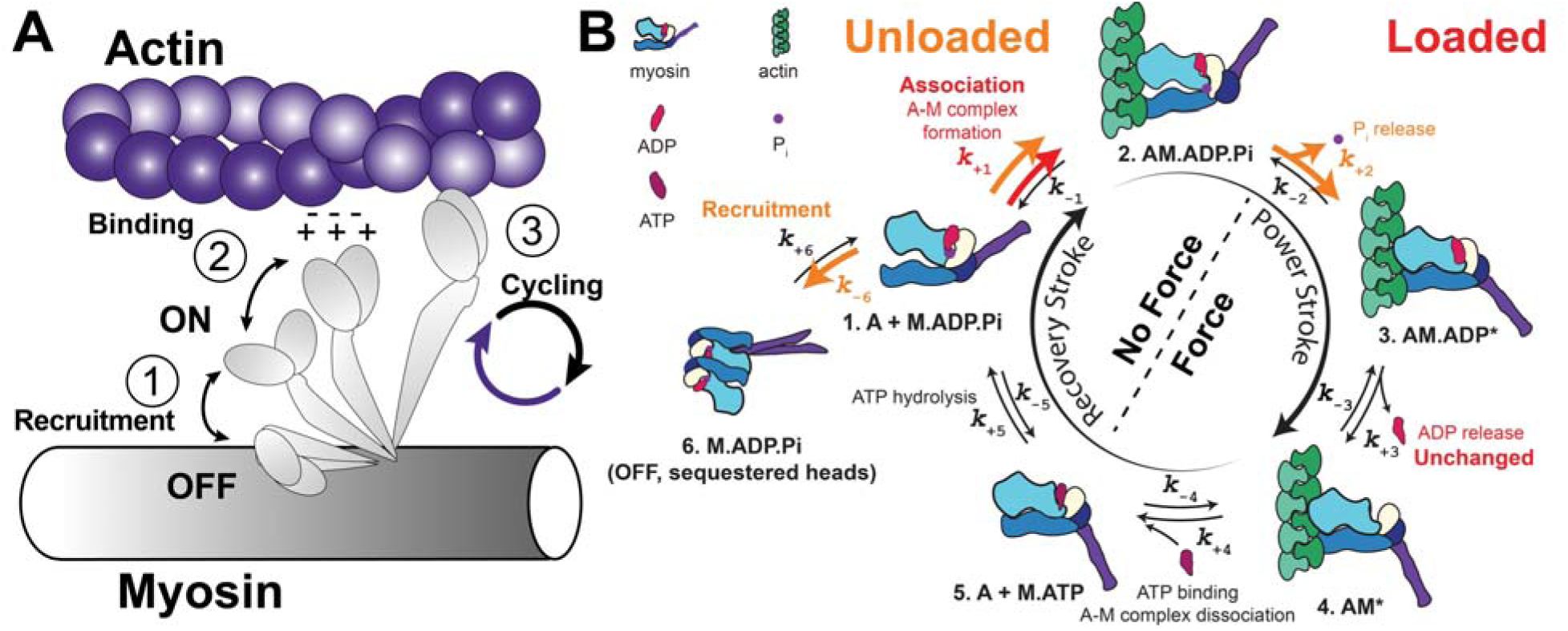
Thick filament regulation and the E525K mutation. (A) Cartoon depiction of myosin and actin highlighting the key steps of thick filament regulation: ① Recruitment of myosin from an OFF to an ON state. ② Actomyosin binding. ③ Myosin cycle rate or ATPase activity. (B) Detailed depiction of the myosin cross-bridge cycle showing all five steps, as well as a sixth myosin recruitment step. Rates of state transition are shown (k_±1-6_) and rates that are changed by the E525K mutation under unloaded conditions are highlighted in orange, rates that are changed under loaded conditions are highlighted in red.

Here we report measurements of contraction and relaxation in intact myofilaments at the myofibril, cell, and tissue level using human induced pluripotent stem cell (hiPSC)-derived cardiomyocytes (CMs) that were engineered to carry the heterozygous E525K mutation. For E525K hiPSC-CMs, force produced by engineered heart tissues (EHTs) and single-cell shortening were decreased when compared with WT controls. However, at the level of isolated myofibrils, E525K hiPSC-CMs generated more force under maximally activated conditions (pCa 4.0) and similar force during sub-maximal calcium activation (pCa 5.6, pCa 5.8). Importantly, there were no differences in contraction or relaxation kinetics from E525K versus WT MYH7 preparations at the EHT, cell, or myofibril levels, suggesting the chemomechanical cycling rate was unaffected. Two results need further study to resolve apparent contradictions: (1) the discrepancy in force generation between cells/tissues and individual myofibrils, and (2) the unaffected kinetics of contraction and relaxation with load, while others have reported altered ATPase activity with the E525K mutation. To resolve issue one, we performed spatially explicit sarcomere modeling to show that our results (increased myofibril isometric force generation and decreased EHT twitch force) can be explained by changing three rates: a decreased rate of myosin recruitment from the OFF/IHM state, and increased rates of actin binding and Pi release from myosin. The reduced rate of myosin recruitment for E525K myosin agrees with recent reports that the mutation increases stability of the IHM conformation, suggesting delayed myosin recruitment out of the OFF state could occur [9,16]. To resolve issue two, we measured ADP release from purified myosin S1 and the effect of elevated ADP on myofibril contractile kinetics. Neither of these was affected by the E525K mutation, supporting the hypothesis that crossbridge cycling kinetics are not altered during isometric contractions.

The results presented here extend our understanding of how the E525K mutation alters myosin function in contracting myofibrils (**Figure 1B**), where the structure of the contractile lattice and strains induced during contraction can affect function. Our results suggest that in human cardiac muscle, reduced myosin recruitment results in hypocontractility for twitch contractions, even with greater myosin affinity for actin and accelerated Pi release. The mutation causes sarcomere/myofibril disorganization that can also contribute to hypocontractility in cardiomyocytes and EHTs. Additionally, hypocontractility is not accompanied by changes in acto-myosin crossbridge cycling kinetics during loaded contraction. These findings underscore the need for multi-scale, multi-disciplinary analysis of protein mutations in cardiomyopathies to completely understand mutation mechanisms.

## Methods

Detailed methods are available in the **Supplemental Materials**.

### Statistical Analysis

Various statistical tests were used to determine significance, depending on the assay, group size, data type, and number of replicates (see figure captions for details). For all tests, a p-value of < 0.05 was considered significant.

### Cell line generation and cardiomyocyte differentiation

hiPSCs were provided by the Allen Institute for Cell Science and differentiated into cardiomyocytes using established protocols to modulate WNT signaling [17]. Advanced maturation was achieved by long-term culture, patterning, and incorporation into engineered tissues, as we have previously described [18,19]. All mechanics and morphology experiments were carried out after 45 days of differentiation and maturation.

### E525K sS1 purification and stopped flow

Recombinant WT and E525K human β-cardiac myosin short S1 constructs were co-expressed with human ventricular essential light chain in differentiated C2C12 myotubes using adenoviral expression and purified by anti-FLAG affinity chromatography followed by TEV cleavage and anion exchange chromatography. Proteins were flash frozen and used for kinetic analyses.

### Molecular, sarcomere, and cell-level modeling

Electrostatic interactions between myosin variants and actin were calculated using *DelPhiForce* [20] after molecular dynamics simulations, as previously described [21,22]. Half-sarcomere simulations were performed using a spatially explicit model incorporating filament elasticity, calcium-regulated thin filament activation, and myosin OFF/ON state transitions [23–25]. Perturbations to OFF/IHM stability, phosphate release, and actin binding were then systematically applied to the WT. Cell-level contraction was simulated with a 2D finite-element model of a single myocyte and its cellular structures.

## Results

### E525K Engineered Heart Tissues and Single Cardiomyocytes are Hypocontractile with Unaltered Contractile Kinetics

WT and heterozygous E525K hiPSC lines were differentiated into cardiomyocytes using established protocols[26,27]. Lactate purification was used to ensure the purity and maturation of all differentiations (**Supplemental Figure 1A**). All experiments were carried out on day 45 of differentiation unless specified otherwise, and all differentiations exhibited homogeneous beating and automaticity in culture. Both cell lines expressed predominantly β-cardiac myosin at the time of the experiments (**Supplemental Figure 1B**).

To determine the effect of the E525K variant on the contractile properties of cardiac muscle tissue, we measured stimulated twitches of engineered heart tissues (EHTs). E525K EHTs had lower isometric force generation when paced at 1 Hz (**Figure 2A and Supplemental Figure 2**). Peak twitch force (**Figure 2B**) was significantly less in E525K (0.751 ± 0.079 mN/mm²) versus WT (1.95 ± 0.127 mN/mm²) EHTs. The tension time integral (**Figure 2C**) was also less in E525K (-1.7e4 ± 3.3e3 %WT*ms) versus WT (1.1e-12 ± 1.6e3 %WT*ms) EHTs. Cross-sectional area (CSA) of the EHTs was comparable between mutant-containing and WT EHTs (**Figure 2D**). Interestingly, the rate of force generation and relaxation in these tissues was unchanged by the E525K variant, as seen in the time to peak and time to baseline measurements (**Figure 2E-F**).

**Figure 2:**
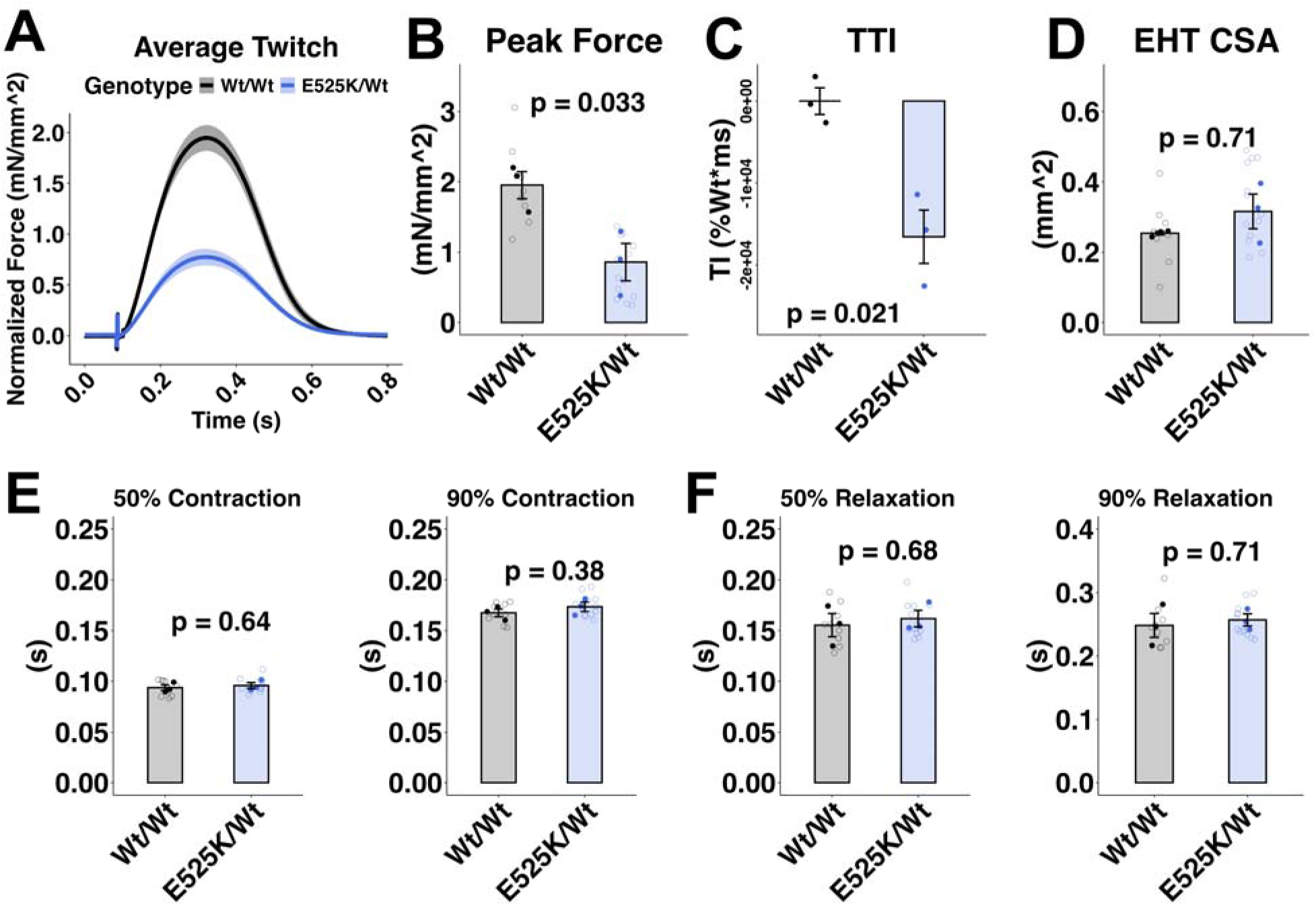
Isometric twitch force is reduced in E525K engineered heart tissues (EHTs) with unchanged kinetics. (A) Average isometric twitch contraction trace (mean ± SEM). Each EHT was paced at 1 Hz and recorded for ∼30 s (Supplemental Figure 2). Individual twitches within each recording were averaged to generate a representative twitch trace per EHT. (B) Peak isometric twitch force. (C) The tension time integral (TTI). (D) EHT cross-sectional area. (E-F) Contraction and relaxation rates. Open circles represent technical replicates (individual EHTs), filled circles represent biological replicates (separate differentiations). All data is represented as the mean +/- SEM. All statistics represent a student’s T-test; N = 3 differentiations, n = 9-13 EHTs per genotype.

EHTs were used here to measure isometric force, which can be challenging to measure in single cardiomyocytes. Single cells were used to address morphological factors that can affect force development in individual cardiomyocytes and EHTs. We measured contraction in live cells using an endogenous eGFP tag on α-actinin for live imaging/tracking of sarcomere movement in WT and E525K cells. To generate elongated and aligned myofibrils and to mature cell morphology, day 45 cardiomyocytes were cultured on lined patterns for one week prior to imaging (see Methods). Both WT and E525K cells contracted along a longitudinal axis when paced at 1 Hz (**Figure 3A-B**). However, E525K cells shortened less (**Figure 3B-C**), as measured by pixel displacement of labeled z-disks: E525K (1.04 ± 0.08 um) versus WT (1.51 ± 0.12 um). The area under the twitch displacement curve revealed a reduced tension time integral (**Figure 3D, Supplemental Figure 3A**), consistent with a hypocontractile phenotype: E525K (-8.65e3 ± 1.45e3 %Wt*ms) versus WT (1.14e-12 ± 1.93e3 %WT*ms). Interestingly, as observed in tissues, there was no change in the rates of contraction or relaxation (**Figure 3E-F and Supplemental Figure 3B**). Together, these results demonstrate weaker contraction in the absence of changes in contractile kinetics.

**Figure 3:**
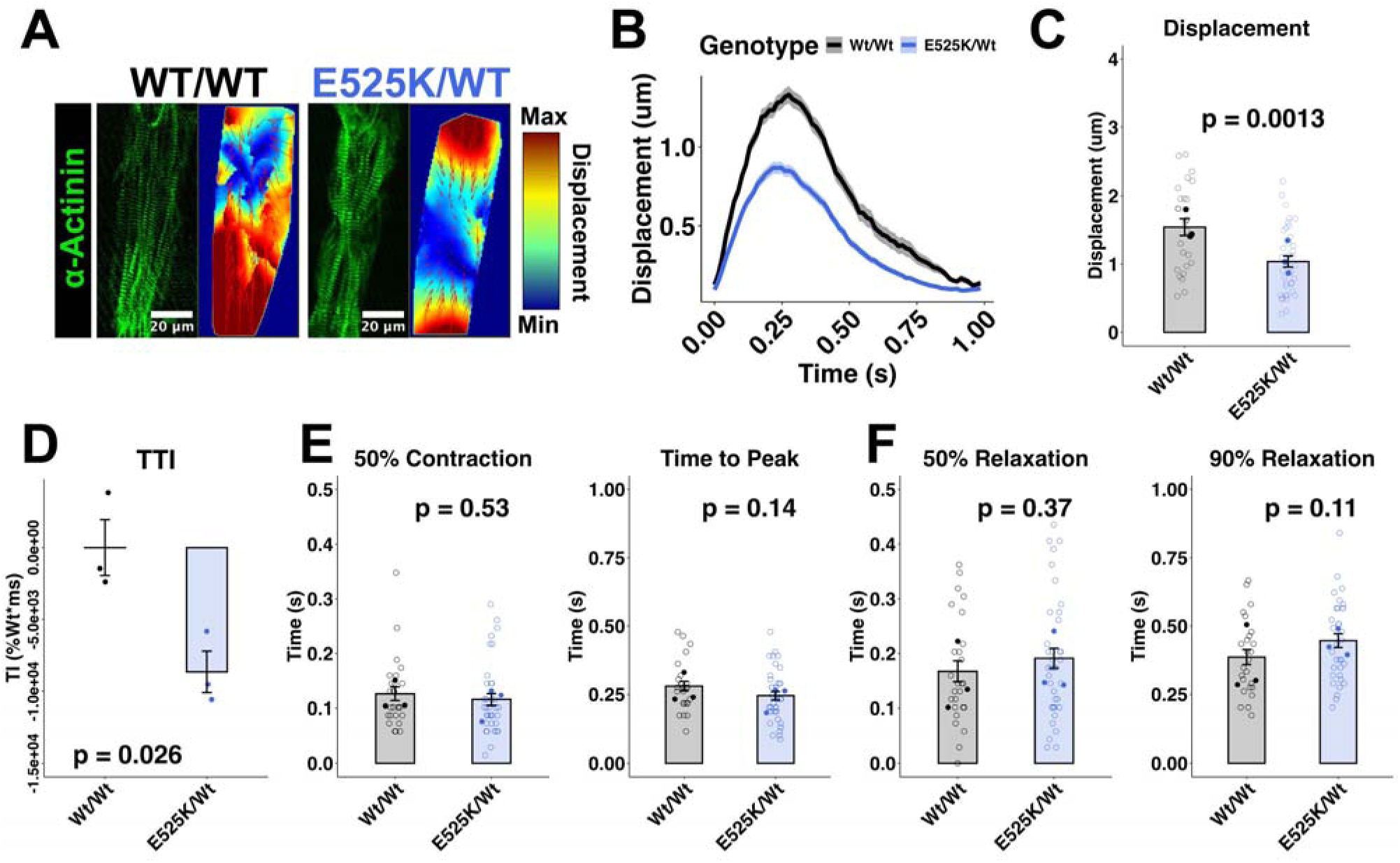
Live cell contraction analysis shows decreased contraction in E525K cardiomyocytes with unchanged kinetics. (A) Patterned day 45 cardiomyocytes with endogenous α-actinin eGFP tag. Representative displacement tracking images are shown. (B) Average trace of displacement over time for live-imaged cardiomyocytes (Mean +/- SEM). (C) Average cell displacement. (D) The tension time integral (TTI) calculated based on the displacement traces. Technical replicates are not shown because paired measurements are required for this analysis. (E-F) Time to peak and time from peak to baseline. Open circles represent technical replicates (individual cells), filled circles represent biological replicates (separate differentiations). Students T-test, N = 3-5 differentiations, n = 91-94 cells per genotype.

### E525K Myofibrils are not Hypocontractile and Have Unchanged Contractile Kinetics

To determine whether a force deficit occurs at the level of individual contractile organelles, myofibrils were isolated from patterned cardiomyocytes and mounted onto glass micro-needles—one of which served as an optically based force transducer [28]. Myofibrils were then activated at maximum and sub-maximum calcium concentrations and relaxed via rapid-solution switching (**Figure 4A**). Unlike in cells and tissues, myofibrils from cells with the heterozygous E525K mutation generated as much force as WT myofibrils when activated with calcium levels similar to those in peak cardiac systole: pCa 5.6 E525K (36.7 ± 2.99 mN/mm²) versus WT (32.5 ± 2.48 mN/mm²); pCa 5.8 E525K (11.3 ± 1.99 mN/mm²) versus WT (8.44 ± 1.01 mN/mm²). Surprisingly, the E525K myofibrils generated significantly more maximally calcium-activated (pCa 4.0) force than WT myofibrils: E525K (51.3 ± 3.45 mN/mm² versus WT (37.0 ± 2.19 mN/mm²; **Figure 4B**). Importantly, the kinetics of activation (*k*_ACT_) and the early, slow (*k*_REL,slow_) and main fast (*k*_REL,fast_) phases of relaxation were unchanged by the mutation at all calcium levels tested (**Figure 4 C-E and Supplemental Figure 4**). These findings demonstrate that the contractile capacity of E525K myofibrils is not compromised under conditions of isometric contraction.

**Figure 4:**
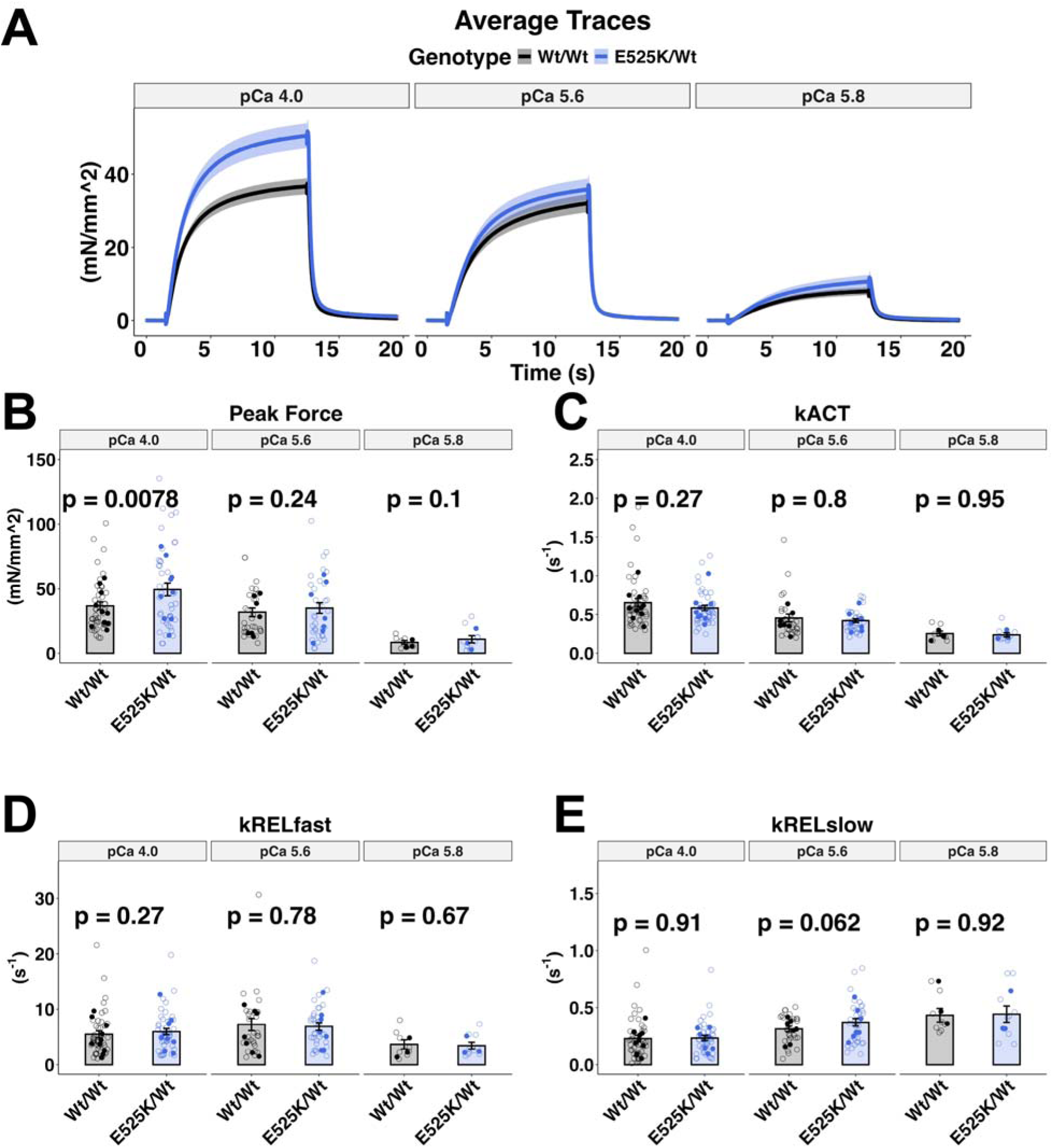
Myofibril mechanics show increased force generation with the E525K mutation under maximal activation conditions with unchanged kinetics. (A) Average traces of normalized force over time (Mean +/- SEM) at maximal (pCa 4.0) and submaximal (pCa 5.6 and pCa 5.8) stimulation. (B) Peak force. (C-E) Kinetics of contraction and relaxation. Open circles represent technical replicates (individual myofibrils), filled circles represent biological replicates (separate differentiations). Linear mixed-effects models test, N = 3-10 differentiations, n = 8-43 myofibrils. Due to the inherent variability of this assay, statistical analysis was calculated based on technical replicates.

The dramatic reduction in force seen in cells and tissues is inconsistent with the preserved or even increased force observed in myofibrils. This disparity is likely not due to differences in calcium handling between the mutant and wild-type cell lines, as calcium transients in patterned and matured hiPSC-CMs showed no difference in amplitude (**Supplemental Figure 5A-B**), baseline (**Supplemental Figure 5C**), or rise and decay kinetics (**Supplemental Figure 5D-E**). To resolve this apparent paradox, we next investigated the cellular morphology and biochemistry of E525K myosin.

### Sarcomere Formation and Organization are Decreased in E525K Cardiomyocytes

Numerous disease-associated variants in thick filament proteins have been reported to influence sarcomere organization in stem cell-derived cardiomyocytes. This can be overcome, at least partially, by external cues such as micro-patterning[26,29,30]. To investigate sarcomere formation in the absence of external morphometric cues, as a general assessment of the variants’ effect on sarcomere/myofibril development, cardiomyocytes were cultured to 38 days on flat surfaces. 7 days before analysis, cells were re-plated to a single cell density. Day 45 cells were then relaxed with blebbistatin and fixed, followed by imaging using the intrinsically expressed α-actinin eGFP reporter (**Figure 5A and Supplemental Figure 6A**). The radial alignment of sarcomeres common to un-patterned hiPSC-CMs was seen in both genotypes.

**Figure 5:**
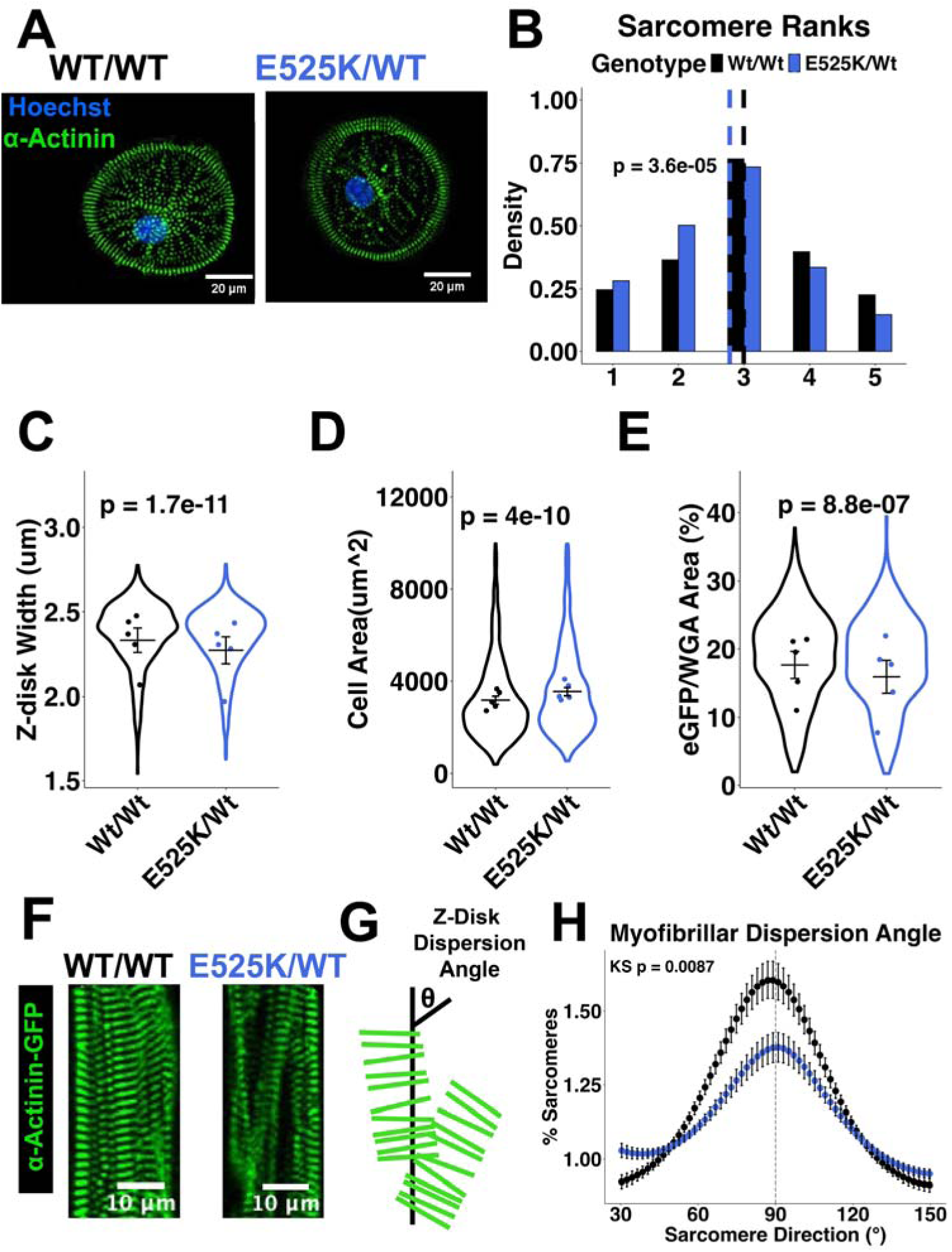
Sarcomere formation and organization is decreased in E525K cardiomyocytes. (A) Representative day 45 un-patterned single cardiomyocytes. (B) Blind ranking of sarcomeres in single cells (scale from 1 “poorly formed” sarcomeres to 5 “well formed” sarcomeres). (C) Z-disk width based on segmentation of endogenous eGFP-tagged α-actinin. (D) Cell area. (E) Percent of cell area filled with sarcomeres/myofibrils. Statistical analysis was performed using a linear or ordinal mixed-effects model with genotype as a fixed effect and differentiation as a random effect to account for non-independence of cells (n = 828-989) within biological replicates (N = 3-5). (F) Representative day 45 patterned single cardiomyocytes. (G) Cartoon depicting the angles measured by dispersion angle analysis. (H) Dispersion angle analysis on whole patterned cells—Kolmogorov-Smirnov Test, N = 5 differentiations.

For organizational analysis, cells were blindly ranked, as previously described [18], based on sarcomere formation and organization (from 1-disorganized to 5-organized; **Figure 5B**). This analysis showed that E525K cardiomyocytes formed fewer and more disorganized sarcomeres. Segmentation of cells and z-disks showed a small but significant decrease in z-disk width: E525K (2.35 ± 0.006 um) versus WT (2.37 ± 0.006 um; **Figure 5C and Supplemental Figure 6C**), and a small but significant increase in cell area: E525K (3530 ± 53.4 um²) versus WT (3144 ± 44.9 um²; **Figure 5D and Supplemental Figure 6B**). E525K cells also had decreased sarcomere content per cell (**Figure 5E**) as measured by percent of cell area containing the eGFP signal: E525K (15.9 ± 2.42 %) versus WT (17.6 ± 1.99 %). Sarcomere content was also quantified in EHTs and was reduced by the E525K mutation (**Supplemental Figure 6D-E**). Since contractile measurements were made on cells after 7 days on patterned surfaces, we next investigated sarcomere organization under these conditions. E525K and WT cells both robustly formed sarcomeres and elongated myofibrils (**Figure 5F**). However, close analysis of the relative alignment of myofibrils and Z-disks within cells revealed differences. To measure how well z-disks were aligned along the axis of contraction, the dispersion angle of each sarcomere image was calculated. A cell with perfectly aligned Z-disks has a dispersion angle (θ) of 90° (to the axis of contraction; **Figure 5G**). This analysis showed that myofibrils and Z-disks from E525K cardiomyocytes were less well aligned with the axis of contraction, even after patterning (**Figure 5H**). This finding prompted us to use a spatially explicit 2D finite-element model of a single myocyte to compare contraction under original versus adjusted sarcomere orientation and structure [31,32]. This showed that while randomly aligned Z-disks do lead to a decrease in cell shortening, misalignment of myofibrils with the axis of contraction leads to an even larger decrease in shortening, even when each Z-disk within a given myofibril remains well aligned with the myofibril axis (**Supplemental Figure 3C**). Importantly, this shows that myofibrillar alignment is a key consideration for force generation at the cell and tissue levels.

Together, these results demonstrate that the E525K variant perturbed sarcomere assembly and structural organization in the absence of external cues, providing a plausible mechanism that can at least partially account for less cardiomyocyte shortening and diminished EHT force generation (**Figures 2 and 3**). Notably, the increased isometric force production measured in isolated myofibrils is only seen at elevated calcium concentrations (**Figure 4**), suggesting that contractile phenotypes are not fully explained by morphological disruption alone. Thus, to further investigate this, we turned to biochemical analyses of purified E525K myosin to measure how the mutation alters enzymatic activity.

### E525K Myosin sS1 has Increased Actin Affinity and ATPase Activity, but No Change in ADP Release Rate

Previous work has shown that E525K myosin sS1 has an increased actin-dependent ATPase activity[9,10]. Moreover, we recently reported that the E525K mutation may increase myosin affinity for actin through disruption of loop 2 conformation[21]. To determine the actin affinity of E525K myosin, we performed stopped-flow kinetic measurements using recombinant human β-myosin sS1 expressed in C2C12 mouse myoblasts. Pyrene-labeled actin–myosin complexes (Apyr·M) were rapidly mixed with ATP to induce myosin dissociation and monitored as an increase in fluorescence to quantify the fraction of myosin bound to actin across a range of sS1 concentrations (**Figure 6A and Supplemental Figure 7A**). These data form a hyperbolic titration curve that can be fit with the quadratic equation to define the dissociation constant of sS1 for actin: *K*_actin_ (**Figure 6B**). The data suggest that E525K myosin sS1 has a greatly increased actin affinity: E525K (<1 nM) versus WT (7.94 ± 0.74 nM). The E525K value falls below the limit of detection for this assay, so the true affinity may be even higher. This increased myosin binding affinity for actin could partially explain the increased force generation by E525K myofibrils during maximal calcium-activated contraction.

**Figure 6:**
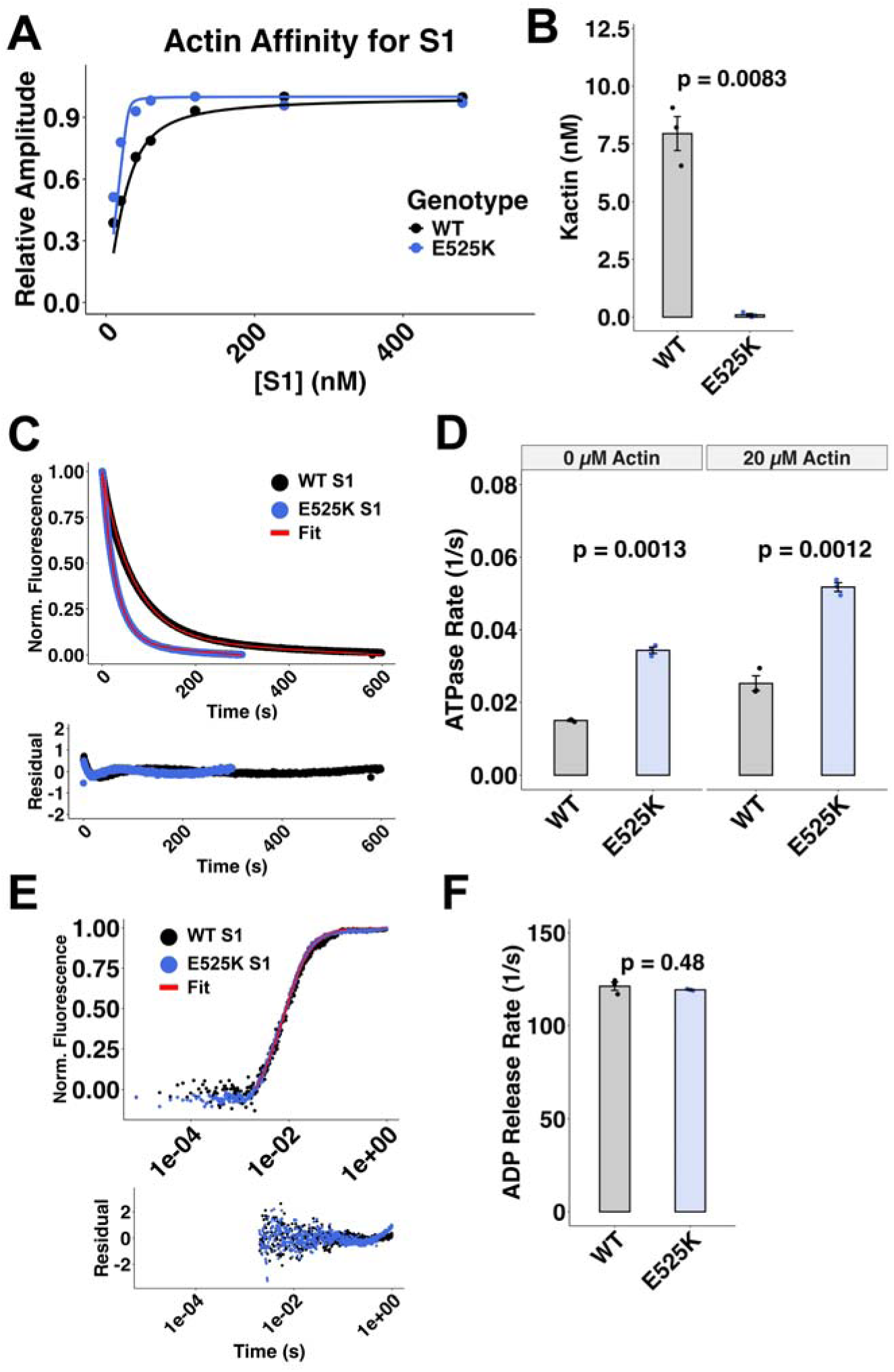
E525K myosin sS1 heads show increased actin affinity, increased ATPase activity, and no change in APD release rate. (A) Stopped flow biochemistry was used to measure the affinity of WT and E525K myosin sS1 (various concentrations) constructs for pyrene-labeled actin (30 nM). The relative decrease in pyrene-actin fluorescent amplitude at various concentrations of myosin sS1 was fit with the standard quadratic equation. An example fit is shown. (B) Dissociation constant or K_actin_ value for each replicate of the pyrene actin assay. (C) mantATP displacement assay was used to measure the ATPase activity of WT and E525K myosin sS1 heads. The decrease in normalized fluorescence was fit with a single order exponential, and residuals are shown. (D) ATPase activity of E525K myosin both in the presence and absence of actin. (E) Rates of ADP release. Change in fluorescence was fit with a two-phase exponential curve. (F) The rates of the fast phase. T-test, N = 3 technical replicates.

Our measures of contraction and relaxation at the EHT, cell, and myofibril levels all suggested no difference in crossbridge cycling kinetics. However, reports by others indicate that isolated E525K myosin sS1 has increased actin-activated ATPase activity when compared with WT myosin[9,10,12]. We confirmed the increased ATPase activity finding using a mantATP displacement assay (**Figure 6C and Supplemental Figure 7B**). In both the absence of actin: E525K (0.034 ± 8.61e-4 s^-1^) versus WT (0.015 ± 1.84e-4 s^-1^), and in the presence of actin: E525K (0.052 ± 0.001 s^-1^) versus WT (0.025 ± 0.002 s^-1^), E525K sS1 heads had a significantly accelerated ATPase rate (**Figure 6D**). This increased ATPase activity has been attributed to increased actin binding and accelerated phosphate release [11], but the release of ADP has not been investigated biochemically. To determine whether ADP release from sS1 is also affected by the E525K mutation, we measured it directly using stopped-flow kinetics. This was accomplished by mixing sS1.ADP bound to pyrene-labeled-actin with a large excess of ATP. Following ADP release, ATP binding to sS1 results in the dissociation of sS1 from actin, which yields an increased pyrene fluorescence. The data were fit with a double exponential (**Figure 6E and Supplemental Figure 7C**), revealing two rates: a fast rate responsible for 80-90% of the total amplitude that reflects sS1 dissociation from actin, and a slow rate that reflects the isomerization that some M.A.ADP complexes must go through to release ADP (**Supplemental Figure 7D and E**)[33]. The fast rate did not differ for E525K sS1: E525K (119 ± 0.28 s^-^1) versus WT (121 ± 2.24 s^-1^; **Figure 6F**), though there was a small but significant increase in the slow rate of ADP release for E525K sS1: E525K (24.4± 0.37 s-1) versus WT (18.8 ± 0.76 s^-1^; **Supplemental Figure 7F**). Notably, the total amplitude of the E525K sS1 trace was greater, consistent with E525K sS1 having an increased binding affinity for actin. Thus, the data suggest there is no substantial change in the ADP release rate from myosin sS1 in solution.

Since force generation and the release of ATP hydrolysis products are strain-dependent processes[14,34,35], we returned to the isolated myofibril assay to determine how elevated ADP affects the rate of the early linear, slow phase of relaxation (*k*_REL,slow_), which reflects the rate of myosin crossbridge detachment[36–38]. Myofibrils were activated and relaxed in the presence and absence of solutions containing 50% ATP and 50% ADP (5 mM total). Under these conditions, there was no difference in *k*_REL,slow_: E525K (0.149 ± 0.0227 s^-1^) versus WT (0.119 ± 0.0161 s^-1^) (**Figure 7B–C**). The relationship between E525K and WT force remained unchanged (**Figure 7A and Supplemental Figure 8**) compared with no ADP (**Figure 4A**): E525K (44.5 ± 5.81 mN/mm²) versus WT (25.3 ± 1.82 mN/mm²). This suggests the rate of crossbridge detachment is unchanged, and the ADP release rate was not affected by the E525K mutation. Additionally, *k*_ACT_ and *k*_REL,fast_ were also not different for E525K myofibrils (**Supplemental Figure 7G–I**). Taken together, our results suggest that during isometric contractions, the E525K mutation does not affect crossbridge cycling dynamics or the kinetics of ATP hydrolysis product release during myofibril contraction.

**Figure 7:**
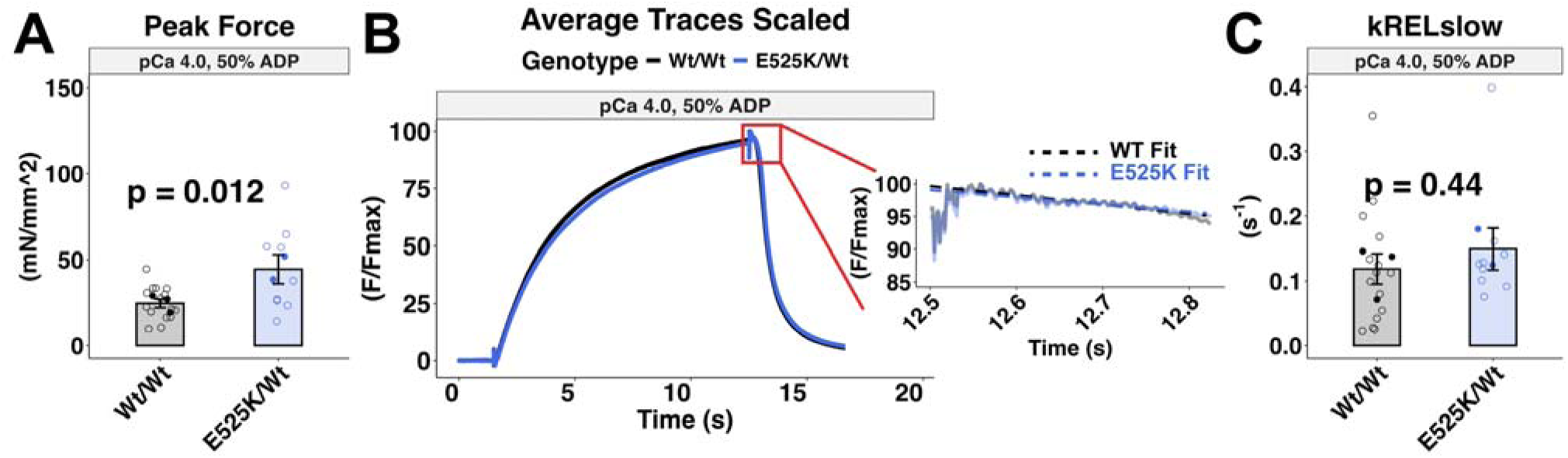
Myofibril mechanics in the presence of ADP show increased force generation with the E525K mutation and unchanged rate of crossbridge detachment. (A) Peak force stimulated at pCa 4.0 in the presence of ADP. 50% of the ATP in both activating (pCa 4.0) and relaxing (pCa 9.0) solutions was replaced with ADP to investigate the biochemistry of ADP release. (B) Scaled average force traces show the kinetics of contraction and relaxation. The zoomed inset graph shows the initial linear slow phase of relaxation as well as the associated linear fit. The scaled traces are shown in translucent lines (gray and light blue), and the linear fits are shown as dashed lines (black and blue). (C) The rate of crossbridge detachment, or *k*_REL,slow_ (slope of the linear fit). Open circles represent technical replicates (individual myofibrils), filled circles represent biological replicates (separate differentiations). Linear mixed-effects models test, N = 2-3 differentiations, n = 9-15 myofibrils. Due to the inherent variability of this assay, statistical analysis was calculated based on technical replicates.

### Electrostatic Force Landscape Alterations in E525K S1 Drive Increased Actin Affinity

Using computational simulations, we recently reported that the E525K mutation increases actomyosin association by modulating the conformational ensemble sampled by loop 2 [21]. Here, we used biochemical approaches to demonstrate that the E525K mutation increases myosin affinity for actin in agreement with a recent report by Bodt *et al.*[11] (**Figure 6 A-B**). To further investigate the mutational influences on actomyosin interactions, we calculated electrostatic force vectors using *DelphiForce*[20,39] across molecular dynamics (MD) trajectories of wild-type (WT) and E525K myosin. As shown in **Figure 8**, myosin molecules (MYH7) were positioned 30 Å from the canonical myosin-binding interface on an actin 5-mer, and the force exerted on each myosin atom by actin was calculated and visualized as vector fields. The force landscapes or vectors of force for each residue in myosin could then be visualized for WT (**Figure 8A**) and E525K (**Figure 8B**) myosin. In both the WT and E525K systems, force vectors at the actin-binding interface are oriented to stabilize association. In the zoomed-in views of the actin-binding surface (**Figure 8A-B, bottom**), direct comparison of WT (black vectors) and E525K (blue vectors) highlights differences in the force felt by loop 2 and a pronounced reversal in the force vector at residue 525 (highlighted red vectors). Notably, the 525-residue undergoes a charge inversion from glutamate (negatively charged) in WT to lysine (positively charged) in the mutant, providing a clear mechanistic basis for the observed electrostatic reorientation.

**Figure 8:**
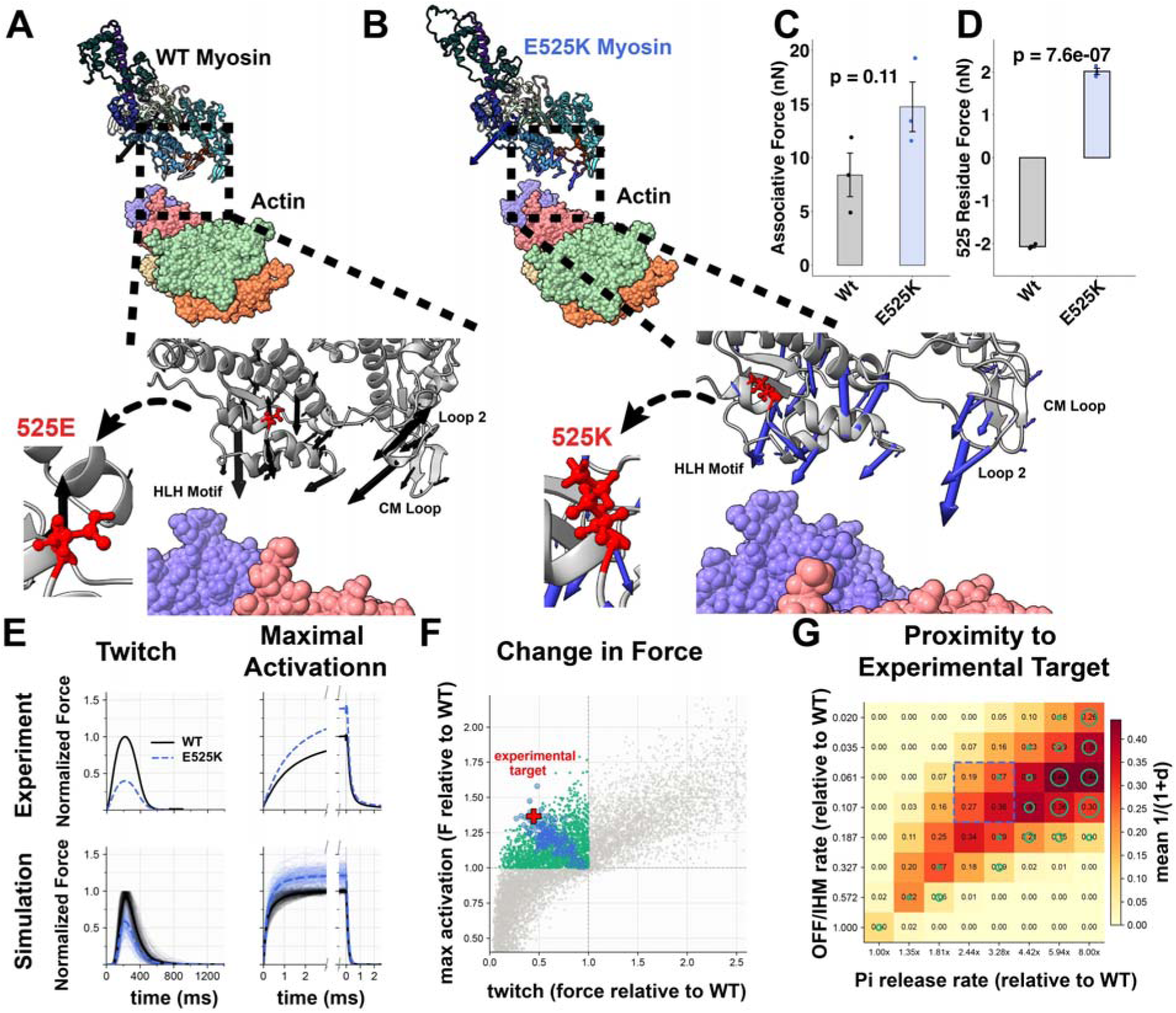
Multiscale modeling elucidates the effect of the E525K mutation on crossbridge formation and force generation. Depictions of WT (A) and E525K (B) myosin (MYH7) in the pre-powerstroke state (ADP.Pi). *DelphiForce* was used to calculate the force vector felt by each myosin atom due to actin, and these vectors are visualized as colored arrows. The large central arrow represents the total force vector for the molecule, and the smaller arrows on each residue represent the force felt by that residue. WT force vectors are visualized in black, and E525K force vectors are visualized in blue. The 525 residues (E/Glu in WT and K/Lys in E525K) are both highlighted in red for ease of visualization. (C) The average associative force across the whole simulation time is shown for each simulation. (D) The average force felt across each simulation by the 525 residue itself (red residues in A and B). T-test, N = 3 separate simulations of each genotype. (E) Experimental (top) and simulated (bottom) traces of twitch and maximal activation. (F) State space relative change after E525K perturbation, three rates were varied: OFF/IHM, Pi Release, and actin binding. (G) Heatmap of proximity of modeled traces to target traces, focusing on OFF/IHM rate and Pi release rate.

To quantify these effects over time, DelphiForce calculations were performed for each frame (1 ns resolution) across three independent 500 ns molecular dynamics (MD) simulations per condition. Substantial variability is observed between replicates, likely reflecting the dynamic conformational behavior of charged structural elements such as loop 2 (**Figure 8C**). Finally, we examined the force experienced by residue 525 over the course of each simulation (**Figure 8D**). Consistent with the structural observations, the direction of the force vector acting on this residue is inverted in the E525K mutant relative to WT. This reversal directly reflects the underlying charge substitution and indicates that residue 525 transitions from contributing to repulsive or neutral interactions in WT to participating in attractive electrostatic interactions in the mutant. Taken together, these results demonstrate that the E525K mutation induces both local and global changes in the electrostatic force landscape governing actomyosin association. The reversal of force direction at residue 525 supports a model in which this mutation enhances actin affinity through altered electrostatic interactions.

### Spatially Explicit Sarcomere Modeling Reconciles the Divergent Effects of E525K on Twitch and Maximal Isometric Force Generation

To investigate the differential impact of E525K on twitch and maximal activation, we used a spatially explicit half-sarcomere model we previously developed (**Supplemental Figure 9**)[24,40–47]. **Figure 8E** shows both twitch and maximal isometric force for both experimental data (from **Figures 2A and 4A**) and simulated data. Each twitch (driven by a simulated calcium transient) and max steady state isometric force (driven by a stepwise change in calcium) simulation is normalized and scaled to WT peak force. WT model traces were generated by fitting the model to twitch and max force traces using a parameter sweep (see Methods). E525K traces were generated by taking the WT parameter set and modulating only rates of OFF/IHM stability (r_x1,6_), actin binding (r_x1,2_), and Pi release (r_x2,3_), as suggested from our experimental data — see Methods and **Supplemental Figure 9** for details. The simulated data shown is drawn from the subset of simulations that responded in a manner consistent with the observed E525K experimental data, e.g. a decrease in twitch force and an increase in max isometric force. Each simulated WT twitch corresponds to a WT max isometric force simulation with the same parameters. Likewise, each simulated E525K trace shown corresponds to a paired WT baseline shown.

For each of the 3776 paired simulations (baseline, perturbation) we calculated the peak twitch and maximum isometric force for both baseline and the E525K perturbation to find the ratio 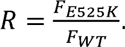 We plotted this ratio for twitch vs maximal isometric force (**Figure 8F**). Each dot represents one of the 3776 (baseline, perturbation) combinations. The upper left quadrant indicates where a perturbation decreased twitch force while simultaneously increasing max isometric force. A black star indicates where the experimental data falls by this measure. While most simulations resulted in both twitch force and max isometric force increasing, a significant portion produced the same change in force as was observed in the experimental data.

We can measure the proximity to the experimental target as 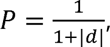 where d is the distance to the experimental target (P = 1 is the best possible score). The proximity values shown in **Figure 8G** are averages across all baselines when the same perturbation is applied to each baseline. The axes indicate the multiplicative change applied to each baseline’s corresponding rate, with the point (1,1) in the bottom left of the plot indicating no perturbation at all. These data show a band in which baselines are perturbed with both increasing OFF state stabilization (decreasing rate) and increasing Pi rate wind up in the correct quadrant. However, outside of this band, one rate or the other dominates. Therefore, in the model, both OFF/IHM stabilization and Pi rate increases are needed to observe the same response as in the experimental data.

While the scatter plot shows that many (baseline, perturbation) pairs end up in the correct quadrant, it is unclear from that plot alone how robust the response is. For example, the plot does not indicate whether this comes from a subset of WT baselines that respond to all possible E525K perturbations in the correct manner, or if every WT baseline responds to just a few E525K perturbations. However, because the heatmap is marginalized across all WT baselines (the plot indicates the average proximity when each perturbation is applied to every baseline), we conclude that the response to the perturbation is robust. Finally, E525K perturbations also included increases to the actin binding rate, and the heatmap shown (**Figure 8G**) also marginalizes over that rate perturbation. While we saw a small effect from increasing the crossbridge binding rate, it was weak and was insufficient alone to generate the experimentally observed behavior. This is consistent with the crossbridge binding rate not being the rate-limiting step, meaning that increasing it does not change the system significantly.

## Discussion

The results presented here represent the first analysis of how E525K myosin impacts myofibril and myocyte contraction, and the morphology of sarcomeres and myofilaments in stem cell-derived cardiomyocytes that are heterozygous for this variant. Myocytes had significantly reduced shortening, and engineered heart tissue (EHT) generated significantly less peak tension (force) and tension over time, consistent with a hypocontractile phenotype [2]. E525K cardiomyocytes had less sarcomere formation and myofibril alignment than WT cardiomyocytes. However, single myofibrils from mutant cells produced a similar force to myofibrils from WT cells at sub-maximal calcium (equivalent to what is experienced during cardiac systole) and had greater maximal force-producing capacity. Finally, while our measurements agree with previous reports of increased ATPase activity for E525K myosin S1, our results in myofibrils, cells, and EHTs all demonstrate that the rates of contraction and relaxation are unchanged. Combined, the data suggest the E525K variant alters force primarily via altered numbers of cycling crossbridges during contractions and not altered crossbridge cycling rates during loaded contractions. This hypothesis was supported by our biochemical and mechanical measurements examining ADP release from myosin, the rate-limiting step during loaded contractions [48,49].

### Multiscale Modeling Elucidates the Mechanistic Basis for Divergent Force Generation in E525K Cells vs Myofibrils

There are several possible explanations for why myofibril isometric force with the E525K mutation was similar at sub-maximal calcium and greater during maximal calcium activation (**Figures 4, 7**), while contraction was reduced in cells (**Figure 3**) and EHTs (**Figure 2**). The lower force in E525K cardiomyocytes and EHTs could result from altered calcium levels during twitch contraction, though we observed no difference in calcium transients in single-cell measurements (**Supplemental Figure 5**). Our morphological analysis suggests that decreased myofibrillar formation and alignment may lead to inefficient force generation along the axis along which force is measured. This was consistent with our modeling of how force is affected by myofibril alignment relative to Z-disk orientation (**Supplemental Figure 3C**). However, this is not a consideration in isolated myofibril force measurements. In myofibril sarcomeres the increased affinity of E525K myosin could facilitate faster and greater crossbridge formation from the pool of readily available myosin. Moreover, the previously published report of actin-accelerated increased Pi release for E525K S1 [11] suggests increased myosin binding for the variant vs WT S1. During thin filament activation by calcium, initial myosin binding can lead to additional crossbridge formation by stabilizing displacement or further displacing tropomyosin from actin binding sites. This is a cooperative positive feedback mechanism in cardiac muscle [50–55]. This study did not directly measure cooperativity, but it likely occurs during the time course of isometric force development in myofibrils. However, it is likely that follow-on myosin binding is less over the ∼200 ms time course of a cardiac twitch, as rapid calcium sequestration back into the sarcoplasmic reticulum counteracts continued myosin binding and further force generation.

The much shorter timescale of twitch vs steady-state isometric contractions could explain why we see hypocontractility in cells and EHTs vs similar or hypercontractility in isolated myofibers, where higher calcium levels are maintained for seconds. Combined biochemical [9–11] and structural biology [9,12] data suggest the E525K mutation promotes a structural conformation that either induces or stabilizes the IHM via increased interaction of the blocked S1 head with S2 in the IHM conformation of myosin[9,10]. The structural bases for this stabilization can be seen in recently published cryoEM structures of the folded-back/IHM state (PDB: 8ACT)[56] and the cardiac myosin filament (PDB: 8G4L)[57]. In both structures, the position of the E525K-residue on the blocked head can be seen near negatively charged residues in the S2 tail (D900 and E903). The E525K mutation could therefore form salt bridges with these residues, stabilizing the IHM. A recent FRET study indicated that the IHM state of the E525K mutant is persistent even in high salt conditions, indicating stronger electrostatic interactions in the E525K mutant stabilizing the IHM[12]. Indeed, EM structures of isolated E525K myosin constructs form more folded/closed conformations *in vitro* [9]. These findings suggest it could take longer for myosin to be recruited from the IHM conformation for E525K cardiomyocytes.

To test whether this proposed mechanism can explain hypocontractile twitches but increased maximal isometric force, we used a spatially explicit half-sarcomere model [24,25,42] fitted to WT twitch and maximal isometric force data (**Figure 8E-G**). We then simulated E525K contractions by changing only three rates (**Supplemental Figure 9**), suggested from existing experimental data: OFF/IHM stability, actin binding, and Pi release. No single perturbation alone was sufficient to recreate the phenotype of decreased twitch force and increased maximal isometric force, because either rate alone dominates, pushing the system out of the experimentally observed regime (**Figure 8G**). The model gives a logical account of why the two measurements diverge. During a twitch, calcium rises and falls quickly, so myosin head recruitment from the OFF/IHM state within this short window is minimal for E525K myosin. When E525K stabilizes the IHM, fewer heads are available for binding to thin filaments before calcium is sequestered, resulting in reduced peak twitch force. Under sustained calcium activation, the time constraint is lessened or eliminated, enabling more myosin recruitment for contractions over time. With increased myosin binding to actin (thin filaments) and faster Pi release, more myosin can be in force-bearing states, thereby preserving force at sub-maximal calcium and increasing maximal calcium-activated force. This idea is bolstered by simulated force-pCa curves, where force is similar for sub-maximal calcium but increased at high calcium (**Supplemental Figure 11**). Notably, our model shows that combining IHM stabilization with acceleration of Pi-release does not alter *k*_ACT_ (Supplemental Table 4). Under this condition, the same rate changes have opposite effects on force depending on the timescale of activation, but as thin filament activation is increased with increasing calcium in myofibril isometric contractions, a greater amount of force can be produced over time. These results support our proposal that the E525K variant slows recruitment of myosin to produce a hypocontractile phenotype in cardiomyocytes and EHTs, limiting the number of myosin heads binding to form cycling crossbridges rather than altering the cycling rates of individual crossbridges. This idea is also supported by the data from us (**Figure 6E-F and Figure 7**) and others [11] that the ADP release rate is unaffected by the E525K variant. The computational analysis with our spatially explicit model only considers the myosin-driven mechanism. Myofibril disarray and misalignment in E525K cells would further reduce twitch force in cells and tissues, as demonstrated by our finite element-based cell-level contraction model (**Supplemental Figure 3C**).

So, how might the E525K mutation reduce the rate of recruitment from the IHM structure? Recent structural work from the Houdusse lab resolved the folded-back/IHM structure of myosin with and without the E525K mutation [58]. The charge change by the switch from E525 to K525 decreases S2 coiled-coil flexibility and abolishes its salt bridge with residue 484. Instead, it permits a salt bridge between residue 525 in the blocked head and the charged residues 900 and 903 in the S2 tail of the free head, providing a plausible mechanism for IHM stabilization. How E525K alters the S1 head in the ON state remains to be examined. Stabilization of the IHM may also affect calcium-mediated thick filament activation, which X-ray diffraction studies show reduces thick filament order and increases the equatorial intensity ratio (I1,1/I1,0), consistent with S1 recruitment toward the thin filaments [59,60]. Though the time course of these effects is unknown, it is plausible that a more stable IHM could slow or reduce myosin recruitment, an effect that could matter more during the brief cardiac twitch rather than the longer time-scale steady-state force generation in myofibrils.

### E525K Myosin Cycling During Contractions

We observed no changes in the rate of contraction or relaxation in any of our mechanical assays, suggesting similar ATP utilization rates for E525K and WT myosin in cardiomyocytes. Previous reports, confirmed in this study, indicate that the E525K mutation increases the intrinsic ATPase activity of myosin S1 heads[9–12]. Recently published work directly measured the actin binding rate of E525K myosin using a pyrene-actin assay and saw an increase in this rate, which was resistant to increased salt (KCl) concentrations [61]. This indicates that electrostatics between myosin and actin regulate this increased rate. Finally, biochemistry and molecular dynamics simulations indicated that Mg and ADP were more flexible in the E525K nucleotide binding pocket [61]. These results suggest that the E525K mutation increases Pi release, which is rate-limiting for ATPase in isolated myosin. Indeed, enhanced computational sampling has recently been used to show that the E525K mutation accelerates ATPase activity [62]. However, these analyses were done in the absence of the organized sarcomere, where steric constraints impose strains (load) in myosin-actin interactions and other sarcomere components during calcium activation. Notably, myosin ATPase activity slows as load increases during contractions, as both Pi release [34] and ADP release [14,48] are strain-dependent processes. Under isometric conditions, ADP release becomes rate-limiting to ATPase. The combination of muscle mechanics and biochemical assays used here suggests that with the addition of load, differences in acto-myosin ATPase activity between WT and E525K myosin are likely negated.

A limitation in our analysis of how the E525K variant affects the mechanical properties of cardiomyocytes is that our hiPSC-CM line is heterozygous for E525K, designed to recapitulate the human condition, so we cannot rule out interactions between WT and E525K myosin in myofibrils from mosaic genotypes. This concern is abrogated somewhat by our protein biochemical analysis of E525K vs WT myosin, which agrees with the literature using pure E525K myosin constructs. Future studies using homozygous E525K stem cell-derived cardiomyocytes or examining myosin heterodimers with one WT and one E525K head could clarify how the mutation influences the various properties of thick and thin filament activation and crossbridge cycling dynamics.

In conclusion, our results support a model in which the E525K variant reduces twitch force generation in cardiomyocytes not through changes in myosin chemomechanical properties, but rather through alterations in the number of myosins that bind to form cycling cross-bridges through alterations in both recruitment and myofibril numbers and organization. Future studies should investigate how this myosin mutation is causing sarcomere disarray. Finally, this study demonstrates the example of a myosin variant that affects crossbridge recruitment in the absence of additional effects on crossbridge cycling dynamics, in contrast to some other myosin variants. This suggests a different mechanism of contractile dysfunction and could affect the response to treatment with myosin-modulating small molecules [63–66]. Additionally, our results suggest that myosin behavior may differ under loaded vs unloaded conditions and across measurement modalities, even for specific myosin mutations, underscoring the need for caution as myosin-modulating therapies for familial forms of HCM and DCM move into the clinic.

## Acknowledgements

This work was made possible by the Allen Institute for Cell Science team, who generated the stem cell lines. The Allen Institute for Cell Science wishes to thank its founder, Paul G. Allen, for his vision, encouragement, and support.

## Supplemental Material

Detailed Methods

Supplemental Figures S1-11

Supplemental Tables T1-4

Supplemental References

