## Supplemental Material for "The E525K β-Myosin Mutation Causes Hypocontractility in Cardiomyocytes Without Altering Crossbridge Cycling"

### Detailed Methods

Cell line generation and cardiomyocyte differentiation

Induced Pluripotent Stem Cells (iPSC) carrying the heterozygous E525K mutation as well as wildtype (WT) controls were kindly provided by the Allen Institute for Cell Science. These cells were generated using established protocols. Briefly, CRISPR Cas9 was used to edit WTC11 iPSC cells with an eGFP tag fused to α-actinin and a heterozygous E525K mutation on MYH7. Cells that went through Cas9 editing but did not receive the E525K mutation were used as wildtype controls for all experiment. Line validation including pluripotency testing and karyotyping was carried out by the Allen Institute for Cell Science.

Cells were differentiated into cardiomyocytes using previous published protocols[1,2]. Briefly, undifferentiated cells (6 × 10^4^) were seeded in Matrigel-coated 12-well dishes in mTeSR supplemented with 10 μM Y-27632 ROCK inhibitor (day -2). After 24 hours media was change dot mTeSR supplemented with 1 μM Chiron 99021. The following day (day 0), media was replaced with RBA media (RPMI supplemented with 0.5 mg/ml bovine serum albumin and 0.214 mg/ml ascorbic acid) and 5 uM Chiron 99021. On day 2, media was changed to RBA with 2 uM WNT C-59. Day 4 media was changed to RBA with no small molecule. On day 6 media was changed RPMI with B-27 supplement. Cells were cultured in RPMI with B-27 with media changes every 2–3 days. To purify cardiomyocytes cells were replated at day 14 and cultured in DEME with no glucose supplemented with 4 mM sodium lactate. Cells were maintained in lactate media for 6 days with media changes every 2 days before being returned to RPMI with B-27. Media was changed every 2-3 days until cells until cells reached the endpoint or protocol described.

Patterning, and maturation

Elongated, mature myofibrils are needed to perform myofibril kinetics experiments. To achieve this from iPSC-CMs the cells were cultured on lined patterns of stamped Matrigel, created using a high-fidelity photoresist double lift-off patterning method. Lined patterns for stamping (1 cm x 1 cm area and 15 um line width) were generated by pouring polydimethylsiloxane (PDMS, Silgard 184, Electron Microscopy Sciences) over a silicon wafer master, cured overnight and then gently peeled off. Ethanol sterilized PDMS stamps were then coated in Matrigel overnight. This Matrigel was then stamped onto an 18 mm cover glass, which was peeled off and then subsequently placed facedown onto a 10 kPa polyacrylamide gel base (0.025 g/mL bis-acrylamide and 0.5 g/mL acrylamide polymerized with 10% APS and TEMED) adhered to plasma treated, glass-bottom 6-well culture dish wells using bind-silane. After hydrating these patterns with phosphate buffered saline (PBS) overnight at 4°C, 300-500,000 iPSC-CMs per well (day 38) were replated onto the patterns in replating media (RPMI, 1X B27 supplement, and 10 mM Y-27632 ROCK inhibitor) and allowed to adhere in a concentrated, 0.4 ml bubble only on top of the patterned area for 2 hours in a 37°C incubator. After 2 hours, each well was topped up with replating media for overnight incubation. 24 hours after replating, media was replaced with 50:50 RPMI:DMEM minus glucose media with 1X B27 supplement to begin increasing the concentration of Ca^2+^ from 0.4 mM to 1.1 mM. After 2 days, Ca^2+^ concentration was further increased from 1.1 mM to 1.45 mM by replacing media with 25:75 RPMI:DMEM minus glucose media with 1X B27 supplement. Cells were maintained in this media for 4 days with one media change after 48 hours (for a total of 7 days on the patterned surface).

EHT Casting, culture, and force measurements

Tissue constructs were generated following established protocols [3,4]. Briefly, rectangular 2% w/v agarose/DPBS casting molds (12 mm length, 4 mm width, ~4 mm depth) were prepared in 24-well plates using 3D-printed spacers. PDMS posts were positioned centrally, upside down, with a 0.5 mm gap between the post tip and mold bottom. Each tissue consisted of 97 μL of a fibrinogen-media solution (87 μL RPMI with B-27 and insulin, 10 μL of 50 mg/mL bovine fibrinogen, Sigma-Aldrich) containing 5 × 10⁵ iPSC-CMs (day 24) and 5 × 10⁴ HS27a human bone marrow stromal cells (ATCC). Immediately before casting, 3 μL of 100 U/mL thrombin (Sigma-Aldrich) was added. The mixtures were incubated at 37 °C for 90 min to allow fibrin gel polymerization around the posts. Gels were then lubricated with media and transferred to fresh EHT media (RPMI with B-27, insulin, and 5 mg/mL aminocaproic acid, Sigma-Aldrich) in a 24-well plate. Media was replaced three times weekly (2 mL/well). Tissues were cultured for 3 weeks and harvested for force measurements at day 45 of differentiation.

For isometric force measurements tissues were removed from posts and mounted on an IonOptix Intact Muscle Chamber System. Tissues were bathed in DMEM/F12 supplemented with CaCl_2_ to a final Ca^2+^ concentration of 1.8 mM. Tissues were field stimulate with 5-10 V and paced at 1 Hz. Paced tissues were allowed to acclimate to the pacing protocol for 5-10 minutes before twitches were recorded for 30 s. Average traces, max twitch force, and contraction/relaxation parameters where calculated using the IonOptix software.

Live cell sarcomere imaging

Patterned day 45 cardiomyocytes were placed in a custom-built live cell imaging chamber on a Nikon A1R Confocal (37°C and 5% CO_2_). Cells were bathed in Normal Tyrode’s buffer (1.8 mM Ca^2+^), stimulated with 5-10 V, and paced at 1 Hz. Live cell recordings of the α-actinin eGFP signal were recorded for 8-10 seconds at a frame rate of 69 FPS. Sarcomere shorting was measured in one relaxed and one contracted frame by hand using ImageJ line scans of two myofibrils form each cell. The angular dispersion of myofibrils and Z-disks was calculated from one relaxed frame of for each cell using the local gradient orientation plugin in ImageJ. Displacement and kinetics of contraction were calculated using the LifeAct software with default settings [5].

Myofibril isolation, and force measurements

For myofibril isolation, day 45 cells on patterned surfaces were directly treated with pCa 9.0 relaxation solution (supplemented with 50 mM Tris, 100 mM KCl, 2 mM MgCl_2_, 1 mM EGTA, 1x protease inhibitor cocktail Sigma, and 1% Triton X-100 Acros Organics) for 10 min on ice. Cells were then washed twice with pCa 9.0 and 1:100 PIC without Triton X-100, gently lifted off the plate with a cell scraper, and then collected off the plate with a pipet. After isolation, iPSC-CM myofibrils were mounted between two glass microprobes, one microprobe was attached to a motor arm and the other served as an optically based force probe, and into a custom-built apparatus with rapid Ca^2+^ solution switching (15°C) to measure myofibrillar contraction and relaxation kinetics at the millisecond timescale. Myofibrils were maximally activated in a pCa 4.0 solution and then subsequently activated in sub-maximal pCa 5.6 and pCa 5.8 solutions.

Immunocytochemistry and immunohistochemistry

Cells were relaxed in blebbistatin (25 mM), then fixed in 4% PFA for 15 minutes. Cells were washed with PBS stained with Wheat Germ Agglutinin (WGA) Alexa Fluor 594 conjugate (ThermoFisher) for 1 hour, with Hoechst 33342 (1:2000) for 10 minutes and mounted in Mowiol 4-88 (Sigma Aldrich). Images were taken on a Nikon AX-R Confocal Microscope. Maximum intensity projection TIFF images of sarcomeres were processed in ImageJ/Fiji using a custom batch macro. Specifically, the WGA channel was used to manually segment the cell area for each cell. This was then used as the ROI to define that cell and measure cell area/shape. For sarcomere analysis in the eGFP/α-actinin channel, each ROI was applied to restrict analysis to the cell area, contrast was enhanced, and intensity thresholding (default dark, 2–65535) was used to segment structures of interest. Thresholded images were converted to binary masks and separated using a watershed algorithm to resolve adjacent objects. Segmented particles were then quantified using the Analyze Particles function (size range 0.50–5.00 µm²), and area, shape, and Feret diameter measurements were reported for each ROI/cell.

Live cell calcium transient imaging

Calcium transients were visualized as previously described [6]. The previously described protocol was adjusted as follows. Patterned day 45 cardiomyocytes were loaded with Calbryte™ 590 AM (AAT Bioquest) to avoid spectral overlap with α-actinin eGFP tag. Cells were consistently loaded for 10 minutes and washed out for 10 minutes. Cells were then placed in a custom live cell imaging chamber (37°C and 5% CO_2_) on a Nikon AX-R Confocal Microscope and bathed in Normal Tyrode’s buffer (1.8 mM Ca^2+^). Cells were stimulated with 10-20 V and paced at 1 Hz. Cells were paced for 30s pacing prior to imaging to allow calcium regulation to reach to steady state. Images were acquired for 9-10 seconds using NSPARC resonant scanning at a frame rate of 50 FPS. NSPARC detector with near 0 noise floor allowed use of lower laser power with high Signal to Noise. ImageJ was used to select a cytosolic region of patterned cells for analysis avoiding the nucleus and endoplasmic/sarcoplasmic reticulum. All cells were analyzed simultaneously using the same settings with the CalTrack automated software in MatLab[6].

E525K sS1 expression and purification

WT and E525K short S1 (sS1, res 1-808) myosin heavy chain (MYH7) was co-expressed with the human ventricular essential light chain (MYL3) carrying an N-terminal FLAG tag cleavable by TEV protease using adenoviruses in murine C2C12 myoblasts (ATCC) differentiated into myotubes; adenoviruses were generated in HEK293T cells (ATCC) using the AdEasy Vector System (Qbiogene Inc, Carlsbad, CA, USA) and purified by cesium chloride gradient centrifugation. MYH7 constructs contained a C-terminal GSG-RGSIDTWV affinity tag. C2C12 cells were grown in 15 cm dishes (~10 dishes per construct) and infected with MYH7 and MYL3 adenoviruses 48 hours post initiation of differentiation into myotubes. Cells were harvested into harvesting buffer (20 mM Imidazole, pH 7.5, 50 mM NaCl, 20 mM MgCl2, 1 mM EDTA, 1 mM EGTA, 10% sucrose, 3 mM ATP, 1 mM DTT, 1 mM PMSF and Roche Protease Inhibitors), 1 ml per dish, 96 hours post infection using a cell scraper, and immediately flash frozen in liquid nitrogen (LN2).

All steps for protein purification were carried out at 4 °C (on ice or in a cold room). Cell pellets were thawed and supplemented with 3 mM ATP, 1 mM DTT, 1 mM PMSF, Roche Protease Inhibitors (1 tablet per 50 mL) and 0.5% Tween-20. Cells were lysed by 50 strokes of a dounce homogenizer and the lysate was incubated for 20-30 mins to allow for the assembly of contaminating endogenous full length (FL) mouse myosin from C2C12 cells into filaments (aided by the low NaCl and high MgCl2 content of the harvesting buffer), which pelleted down in the subsequent spin at 30,000 rpm for 30 mins at 4 °C. The clarified supernatant was incubated with anti-FLAG resin for 1-2 hours to capture β-cardiac sS1 protein, followed by washing away non-specifically bound proteins by centrifugation using wash buffer (20 mM Imidazole pH 7.5 containing 150 mM NaCl, 5 mM MgCl2, 1 mM EDTA, 1 mM EGTA, 1 mM DTT, 1 mM PMSF, 3 mM ATP, 10% sucrose and Roche protease inhibitors). sS1 was then dissociated from the resin by overnight incubation with TEV protease, cleaving the FLAG tag on ELC. The next morning dissociated sS1 in the supernatant was collected by spinning down the resin and the protein was further purified (to remove TEV protease and endogenous mouse FL myosin) by anion exchange chromatography using a 1 mL HiTrap Q HP column (Cytiva) attached to an ÄKTA FPLC system. Myosin typically elutes at ~250 mM NaCl; fractions were pooled based on SDS-PAGE analysis, aliquoted, and flash frozen in LN2 for stopped flow kinetic experiments.

Stopped flow

Constructs of cardiac myosin were recombinantly expressed and purified from the mouse myoblast C2C12 cell line. This allowed for comparison of pure E525K myosin with pure WT myosin unlike in hiPSC-CMs where the mutation was heterozygous. Stopped-flow kinetics were performed at 20°C using a Hi-Tech ‘KinetAsyst’ stopped-flow (SF-61DX2, TgK Scientific, UK). Measurements were performed at 120 mM KCl, 5 mM MgCl_2_, 20 mM MOPS, 1 mM DTT, pH 7.0. For the actin affinity assay pyrene fluorescence was measured by excitation at 365 nm and emission through a GG 400 nm cutoff filter. Actin affinity (K_actin_) was assessed via ATP-induced dissociation of pyrene-actin and sS1 complex. Briefly, 30 nM pyrene labelled actin stabilized by phalloidin was premixed with various sS1 concentrations (5 - 500 nM) and then rapidly mixed with 20 uM ATP. An average trace of at least eight individual traces was generated for each condition and fit with a double exponential. The relative amplitude of fluorescence increase (pyrene-actin dissociation from sS1) was plotted vs. sS1 concentration and fitted with the classical quadratic equation to define the dissociation constant of sS1 for actin (K_actin_) as described previously[7]. For the ATPase activity or single turnover analysis 0.15 μM sS1 was incubated with 2 μM mantATP before rapidly mixing with an excess (250 μM) of MgATP. An average trace of eight to ten individual traces was generated for each condition and fit with a single exponential. The kinetics of florescence decay describe the ATPase rate[7]. ADP release rate was measured as previously described[8]. Briefly, 0.33 μM sS1 + 0.5 μM pyrene-actin was pre-incubated with 80 μM ADP before rapidly mixing with a large excess (1.5-2.5 mM) of MgATP. Traces were fit with a double exponential. The kinetics of the fast phase (k_obs,fast_) describes the ADP release rate.

Myosin Separation Gels

Cell pellets were flash frozen in liquid nitrogen and stored at -80°C until ready to use. Cell pellets were resuspended in ~3x volumes myofibril lysis buffer (5% SDS, 50 mM Tris (pH 8.5), 0.75% sodium deoxycholate, and 1x protease inhibitor cocktail). Sarcomeres and proteins were denatured by boiling samples for 10 minutes. Concentration was measured by blotting for total protein and adjusting loading as needed. Myosin separation gels were made by combining a stacking gel (2.95% Acrylamide with 10% glycerol at pH 8.8) and resolving gel (6% Acrylamide with 10% glycerol at pH 8.8). Separation gels were run for 2 hours and 15 min at 32 mA in a cold room (4°C). Gels were stained for total protein using One-Step Lumitein™ Protein Gel Stain, 1X following manufacturer instructions. Gels were imaged on a BioRad ChemiDoc using SYPRO Ruby protein stain setting.

DelPhiForce Calculations

Electrostatic binding forces between myosin loop 2 variants and actin were computed using *DelPhiForce*[9,10]. Representative frames (1 frame/ns) from molecular dynamics simulations were aligned to a reference human cardiac actin–tropomyosin–myosin complex (PDB: 8EFH) using the MatchMaker tool in UCSF Chimera. An equilibrated actin pentamer derived from this structure was positioned such that the central monomer interacted with myosin. To approximate a weakly bound configuration, myosin structures were aligned only via the helix–loop–helix (HLH) motif (residues 515–550 and 530–553), consistent with previously described weak-binding states (PDB: 8R9V). Myosin was then translated 30 Å away from actin along the normal vector of the interaction plane defined by steric clashes, generating a pre-binding configuration. Structures were prepared using *DelPhiPka*[11], and electrostatic forces were calculated by solving the Poisson–Boltzmann equation with *DelPhiForce* using default parameters. This approach isolates electrostatic contributions to actomyosin interactions and enables comparison of residue-dependent effects on binding.

Half Sarcomere Modeling

Our model follows from the lineage of models described in [12–20]. This muscle model is spatially explicit, meaning each thick and thin filament consists of a chain of crossbridges and actin binding sites, respectively, coupled axially elastically together. These thick and thin filaments are arranged into a 3D hexagonal lattice. Titin, modeled as an exponential spring, connects the ends of thick filaments to the z-disk at a point co-located with where the thin filaments attach to the z-disk, as in [17,19].

Crossbridges are formed of a two-spring system, a linear spring comprising the globular domain, and a torsional spring comprising the converter domain. The spring stiffnesses (*k_r_*, *k_theta_*) and equilibrium position (*r_r_*, *r_theta_*) depend on the chemo-mechanical state of the crossbridge. In our model, crossbridges are modeled by a six-state process, including an unbound ADP state (ON/DRX), loosely bound pre-power stroke, post-power stroke, rigor-like, and an unbound ATP state. The ON/DRX state can also transition into (and back out of) the OFF/SRX state a state of very low ATP turnover. In this model, the OFF to ON transition is calcium modulated, following the formulation in [21].

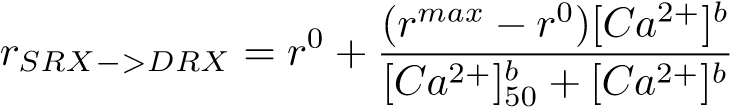

Because the model is spatially explicit, the chemo-mechanical state, local axial displacement, and location within the sarcomere are tracked for each individual crossbridge.

Our simulations require prescribing sarcomere length and calcium for the time course of a simulation. Sarcomere lengths were set at 2200 nm for all simulations, and pCa for maximal isometric activation was set to 4. For twitches, calcium transients came from ratio-metric time-series data that corresponded to EHT twitches (**Figure 2A**). To approximate diastolic and systolic calcium during a cardiac twitch we scaled the trace to have a minimum pCa of 9, and a max pCa of 5.8.

Because of how many parameters are present in the model, the model is in general under-determined when comparing to specific target data, meaning many different combinations of parameters might result in indistinguishable data. Similarly, such parameter combinations which produce indistinguishable simulations might respond differently to the same perturbation. Therefore, rather than simulate a single WT candidate and perturbation, we select many random WT candidates and a broad range of perturbations based on the hypothesis that the E525K variant affects OFF/SRX stabilization, *P_i_* release, and actin binding affinity. We then applied a range of perturbations to each WT candidate. In this way we attempt to characterize the response to various candidate E525K perturbation of the region in parameter space that most reasonably corresponds to the WT data. We started by choosing parameters to vary to characterize the model’s behavior. We simulated 1024 pairs of twitches and maximally activated protocols, where each of the 1024 simulations has random parameters, centered on values consistent with literature and previous models. We generated 1024 candidate parameter sets using Latin Hypercube Sampling (LHS)[22]. Each of the free parameters was divided into 1024 equal probability bins, and then exactly one sample was drawn per bin per parameter, then samples were combined into 1024 complete parameter sets by random permutation across parameters. This is a standard method in randomly sampling high dimensional spaces since it guarantees uniform marginal (but not joint) coverage of the region of interest, for a fixed number of simulations, compared to random sampling or grid sampling, which can jointly sample a region, but for exponentially more simulation cost.

We then rank the 1024 pairs of twitches and max isometric protocols by their similarity to the WT data. We chose twitch peak (*F_max,twitch_*, twitch time to peak (*t_peak,twitch_*), and time to 95% max force in maximal isometric force (*t*_95_*,pCa*_4_). For each metric, *x*, we calculate
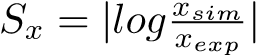
. By taking the ratio, we make all the different metrics unit-less, and by taking the absolute magnitude of the log, we penalize overshoot and undershoot equally. A score of 0 would indicate perfect agreement since log(1) = 0. Of the 1024 parameter combinations initially simulated, we eliminated any that failed to produce at least 20 kPa (half of the WT max isometric force), and then ranked the remaining by their similarity score S. We kept the best quartile of parameter combinations, which formed our WT candidates. These represent plausible WT parameters sets. We were left with 118 WT baselines from this process. For the E525K perturbations, we decreased *r_max_*, increased rate of phosphate release (*r_Pi_*), increased crossbridge binding to actin (*r_xb_*). We again chose random parameter perturbations using LHS. This resulted in 32 perturbations, and each perturbation was applied to each of the WT candidates. Therefore, each of the 118 WT baselines received 32 separate perturbations for 3776 simulations each for twitch and max isometric force protocols.

Modeling differences in myofibril and sarcomere structures

To quantify the spatial dependencies of myofibril and sarcomere structures, we employed a spatially explicit, finite-element mechanical model of a single myocyte. This framework extended previous spatially explicit models of cell-matrix structures [23,24] by going one level deeper to an explicit 2D representation of subcellular structures. These included sarcomeres, Z-lines, connecting units between Z-lines, the cytoskeleton, and nuclei. Z-lines and connecting units were treated as significantly stiffer (15 times in magnitude compared to the sarcomeres), while the cytoskeleton was treated as softer (1/4 in magnitude compared to the sarcomeres). Sarcomeres, as the primary contractile units in the model, were assigned active tension time-dependent transients.

To distinguish myofibril orientation from sarcomere structure/Z-disk orientation, we compared three scenarios of active contraction. An axis of contraction was assigned to each sarcomere: (1) In the original/aligned configuration, this axis followed the baseline orientation of the sarcomere. (2) Random alignment per-sarcomere of Z-disks, within a range relative to the axis of contraction. (3) Random alignment per myofibril, within a range relative to the axis of contraction. Cell shortening over time was tracked as output.

### Supplemental Figures

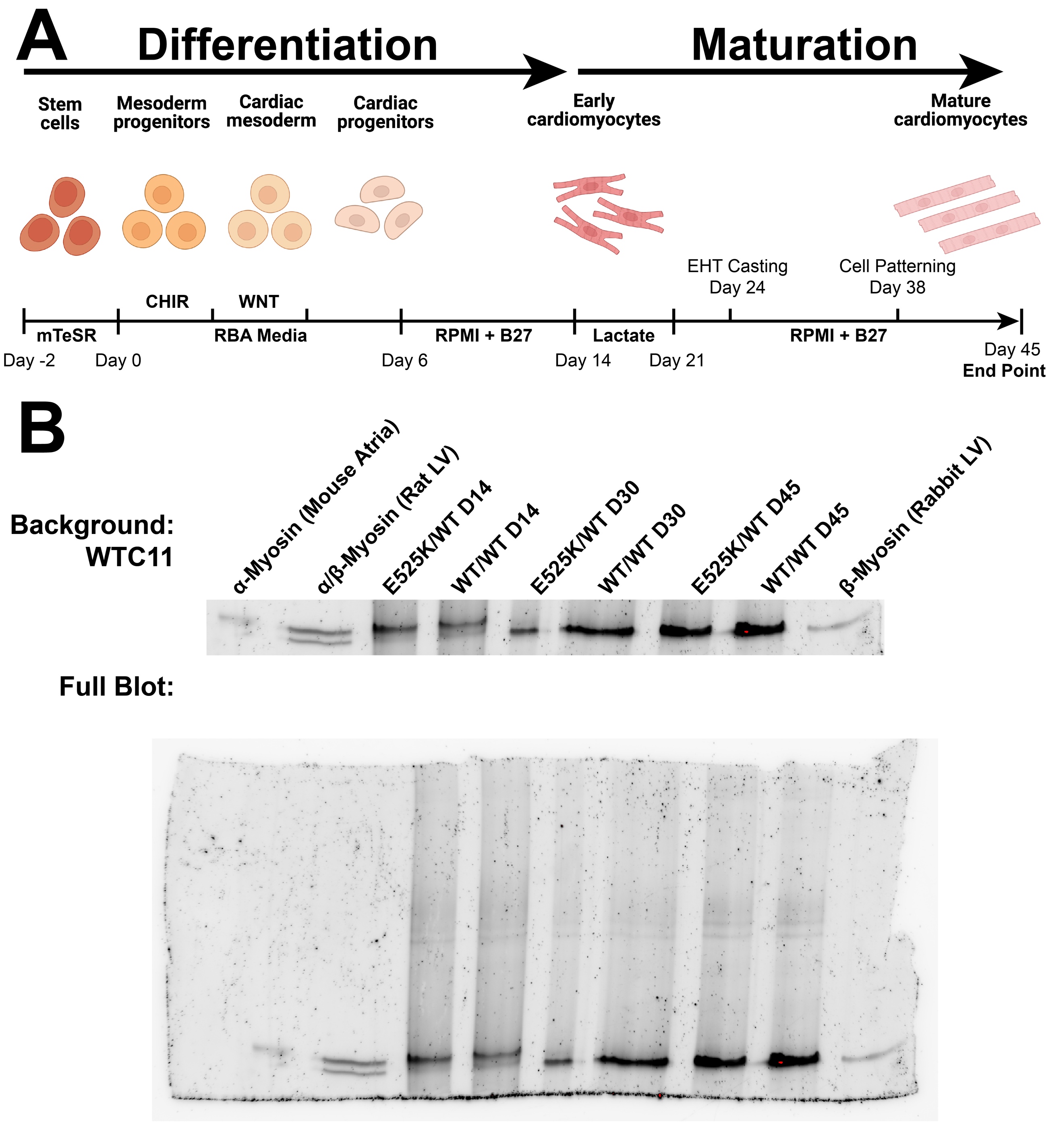

Supplemental Figure 1: Differentiation and maturation of iPSC derived cardiomyocytes

(A) Protocol for the differentiation and maturation of stem cell derived cardiomyocytes from iPSCs. All cell lines underwent the same differentiation and maturation protocol. Differentiated cardiomyocytes were purified by lactate selection from day 15 to day 21. Engineered heart tissues (EHTs) were cast on day 24 and cultured for 21 days on posts. Cells were seeded onto patterns on day 38 and cultured for 7 days on patterns. All end point assays were carried out on day 45 of differentiation. (B) The expression of β-myosin was confirmed using a myosin blot. Day 14 (pre-lactate selection) cardiomyocytes of both genotypes expressed a mixture of α- and β-myosin. After purification (day 30 and day 45) all genotypes expressed only β-myosin. Rabbit left ventricular (LV) heart tissue was used as positive controls β-myosin.

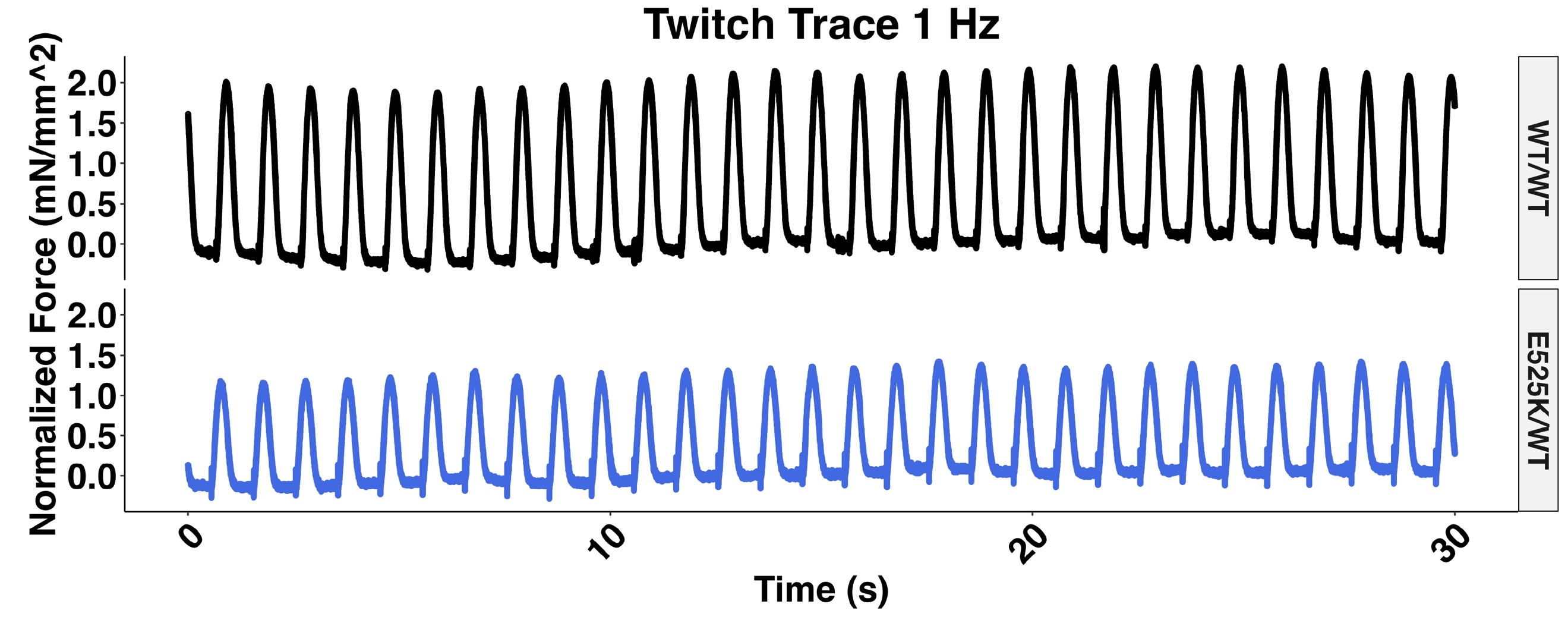

Supplemental Figure 2: Example EHT traces

Example traces from WT and E525K EHTs paced at 1 Hz. Traces were recorded for 30 seconds, and average traces were generated for each EHT using IonOptix software.

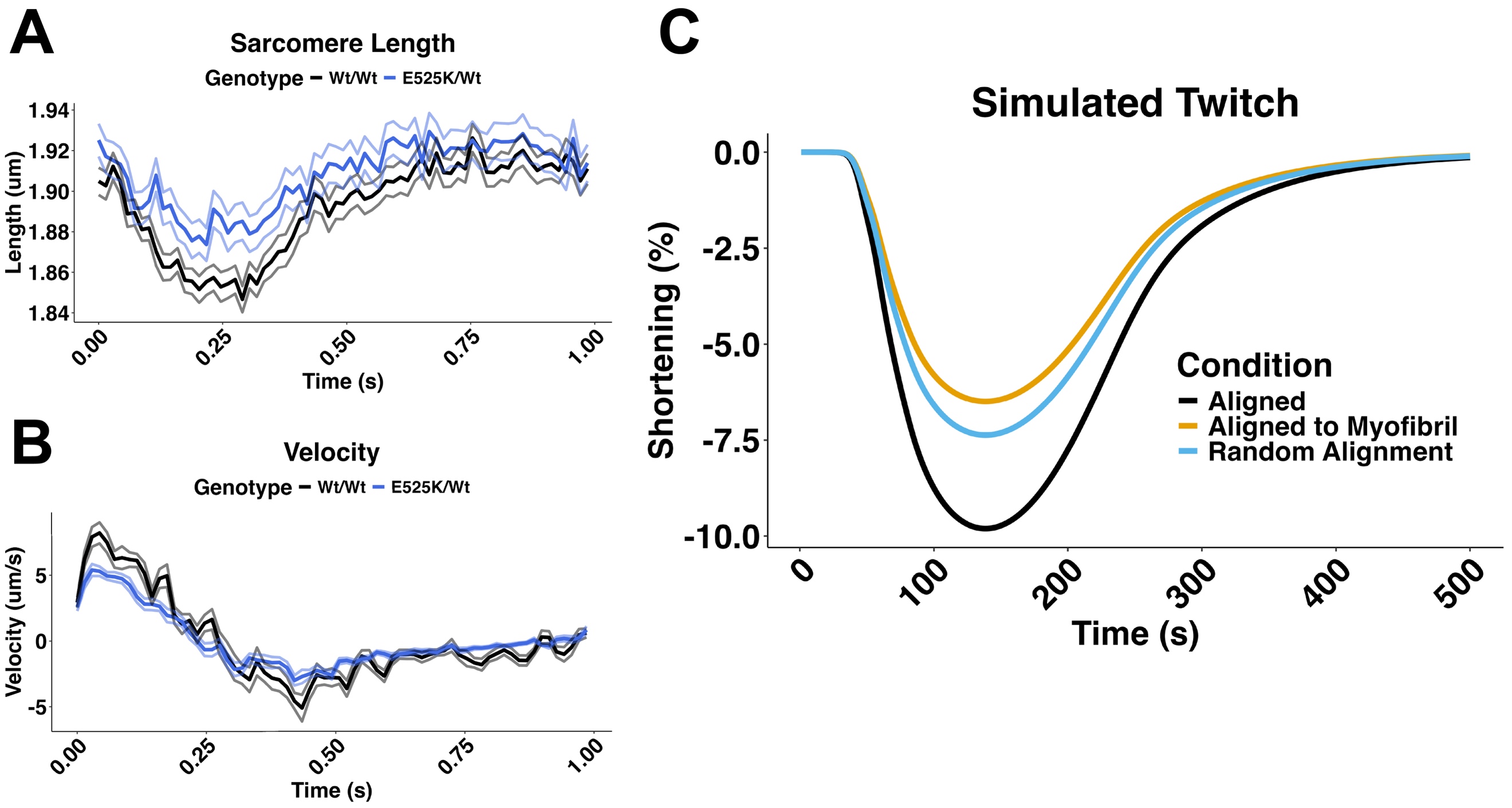

Supplemental Figure 3: Single-cell imaging measurements

(A) Analysis of sarcomere length over time in live imaged iPSC-CMs (B) Analysis of sarcomere velocity over time in live imaged iPSC-CMs. C) 2D finite-element comparison of contraction under original versus adjusted sarcomere orientation and structure, with cell shortening tracked over time for each configuration.

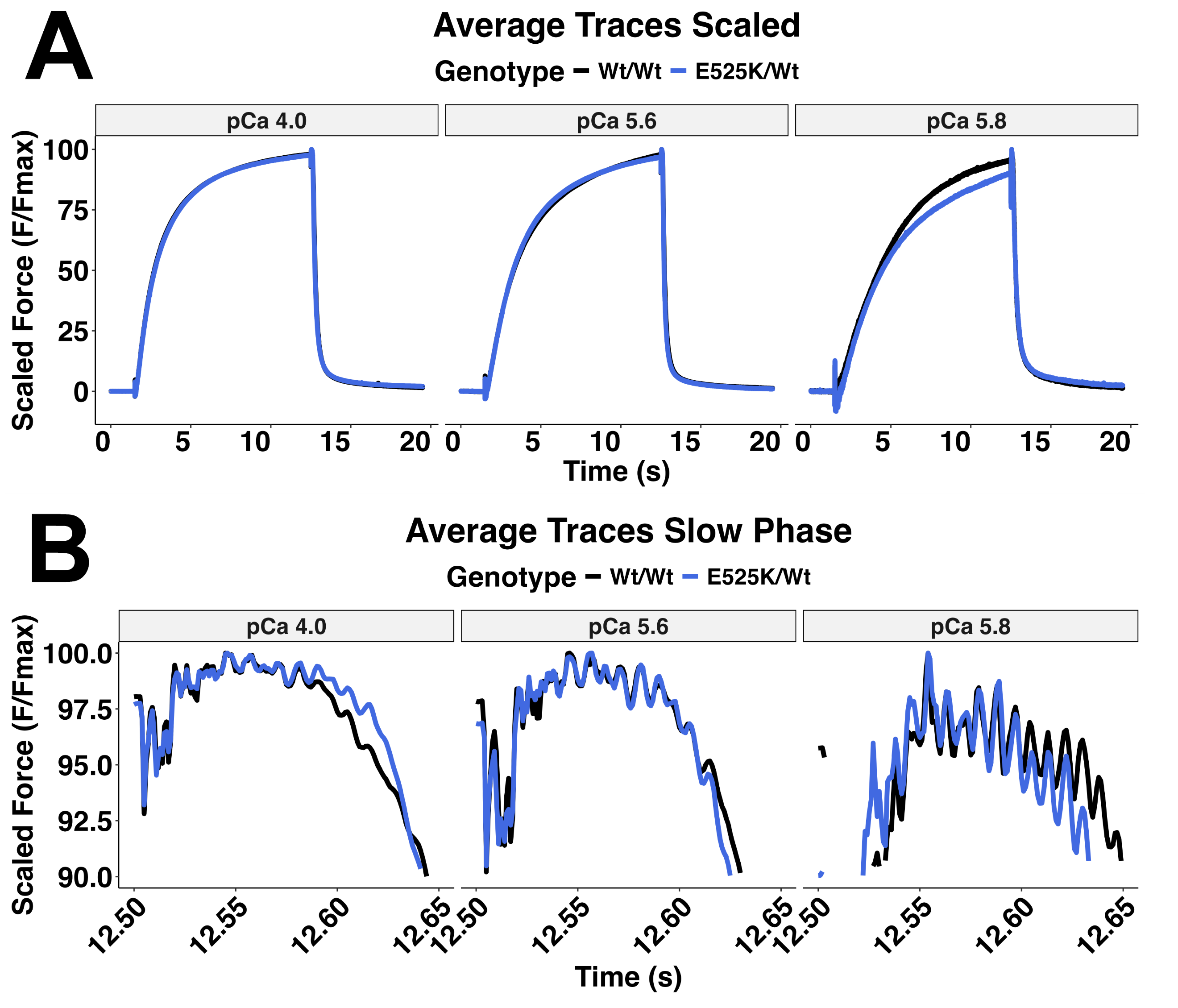

Supplemental Figure 4: Scaled myofibril traces

Whole myofibril traces scaled to max force to compare the shape of the traces for WT and E525K myofibrils (A) and partial traces zoomed in to show the slow phase of relaxation (B). Here, the slow phase of relaxation is shown from 12.5 s (when the myofibril is switched to a relaxing solution) to 12.65. For experimental analysis, the slow phase was determined by manually identifying the inflection point between the linear slow phase and the exponential fast phase for each myofibril trace.

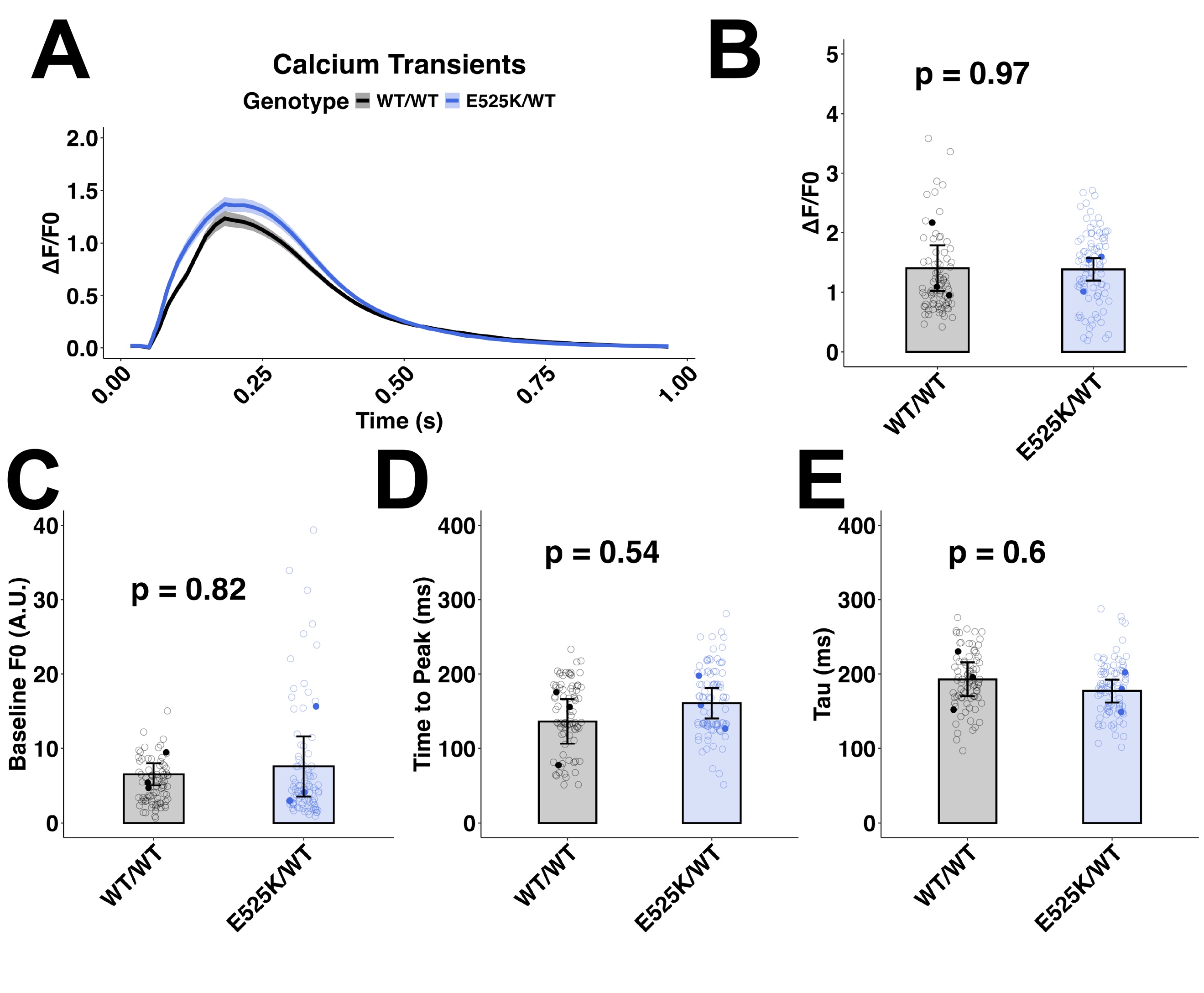

Supplemental Figure 5: Calcium transients are unchanged in E525K cells

(A) Average calcium transient from day 45 patterned cardiomyocytes paced at 1 Hz (Mean +/- SEM). (B) Transient amplitude was unchanged. (C) Baseline fluorescence was unchanged. (D) Time to peak was Unchanged. (E) Rate of calcium return to baseline Tau was unchanged. T-test, N = 3 differentiation, n = 82-92 cells.

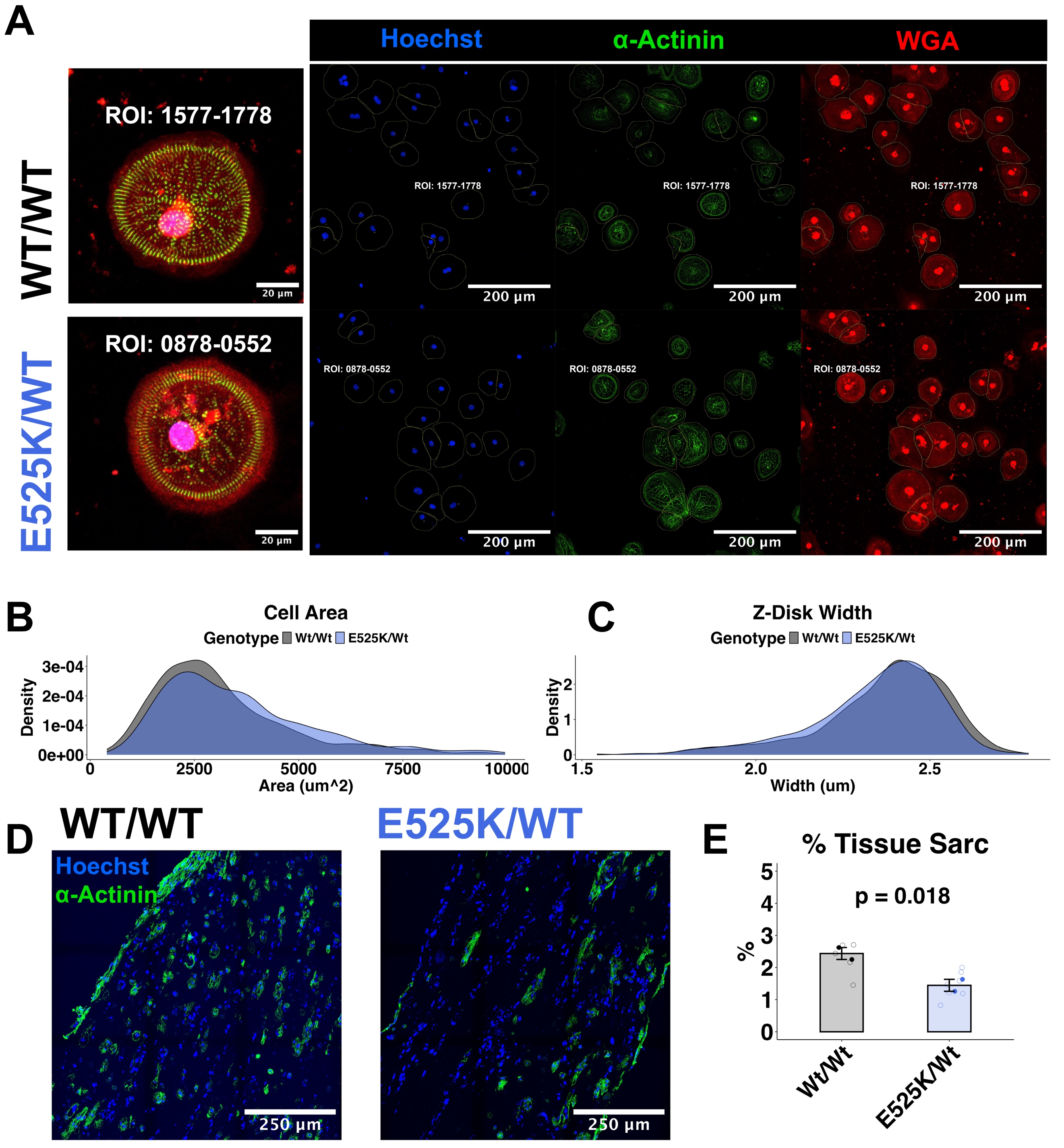

Supplemental Figure 6: Morphology of single cells and engineered heart tissues

(A) Left: example WT and E525K cells. Cells were re-plated to a single-cell density 7 days before fixation on day 45 of differentiation. Cells were stained with Wheat Germ Agglutinin (WGA) Alexa Fluor 594 (red) and Hoechst 33342 (blue). An endogenous eGFP tag on α-actinin was used to visualize z-disks. Right: unmerged images are shown. The representative cell/ROI for each genotype is indicated by the inset ROI number. (B) Density plot of cell area based on segmentation of WGA stain. (C) Z-disk width calculated as the max Feret diameter of segmented Z-disks. N = 3 differentiation, n = 828-989 cells per genotype. (D) Example sections from EHTs are shown, with nuclei (Hoechst 33342, blue) and z-disks (endogenous eGFP tag on α-actinin, green) visible. (E) quantification of the percent of tissue area made up of sarcomeres. Segmentation of the z-disk/eGFP signal was used as a proxy for sarcomere area.

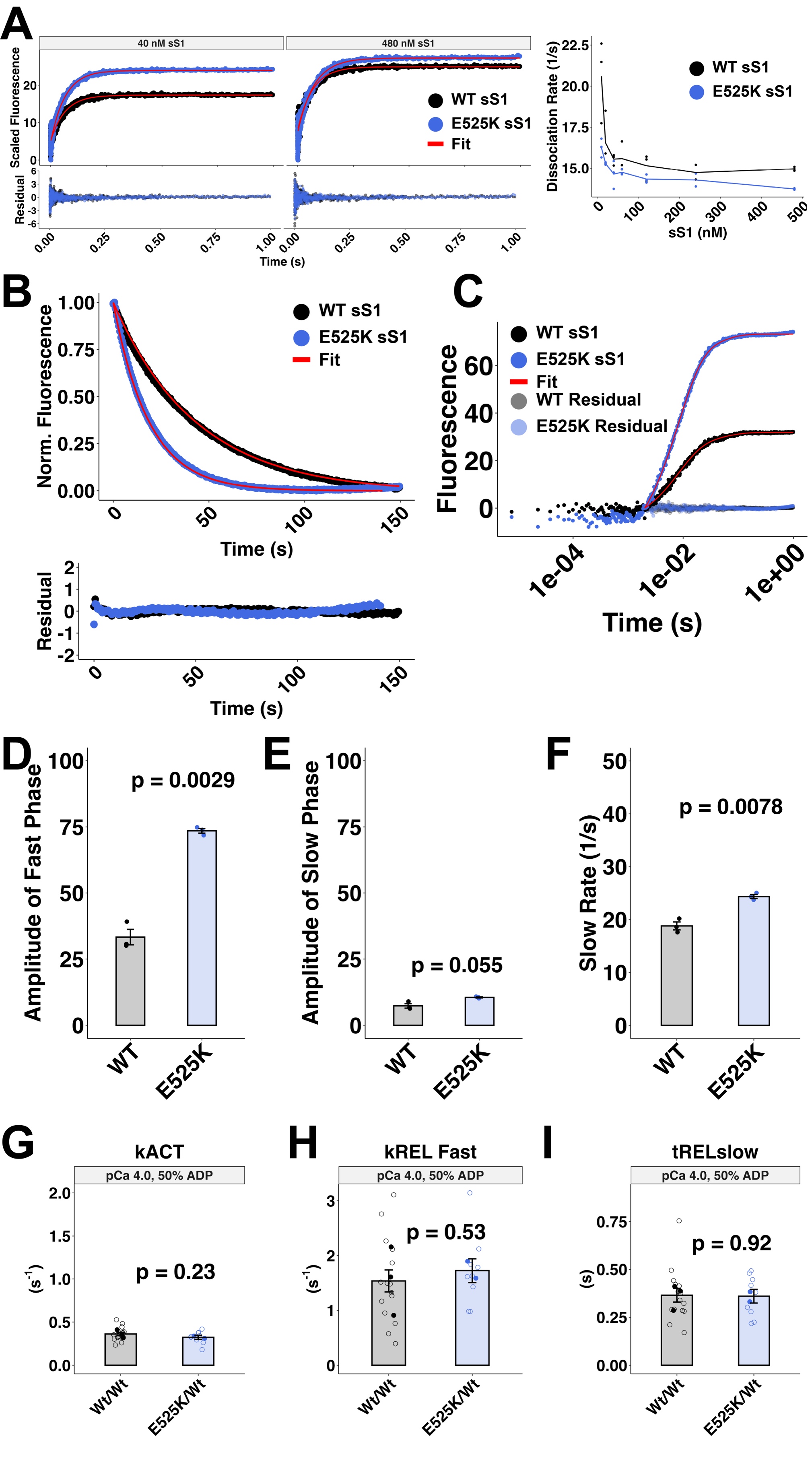

Supplemental Figure 7: Additional stopped flow and myofibril data

(A) Example pyrene-actin displacement traces used to calculate sS1 actin binding affinity. (B) Example mantATP displacement traces for WT and E525K sS1 myosin in the presence of 20 uM Actin. (C) Example unscaled traces of ADP displacement assay for WT and E525K sS1 myosin. (D-F) Additional ADP displacement parameters. (G-I) Rate parameters for myofibrils in the presence of 50% ADP and 50% ATP.

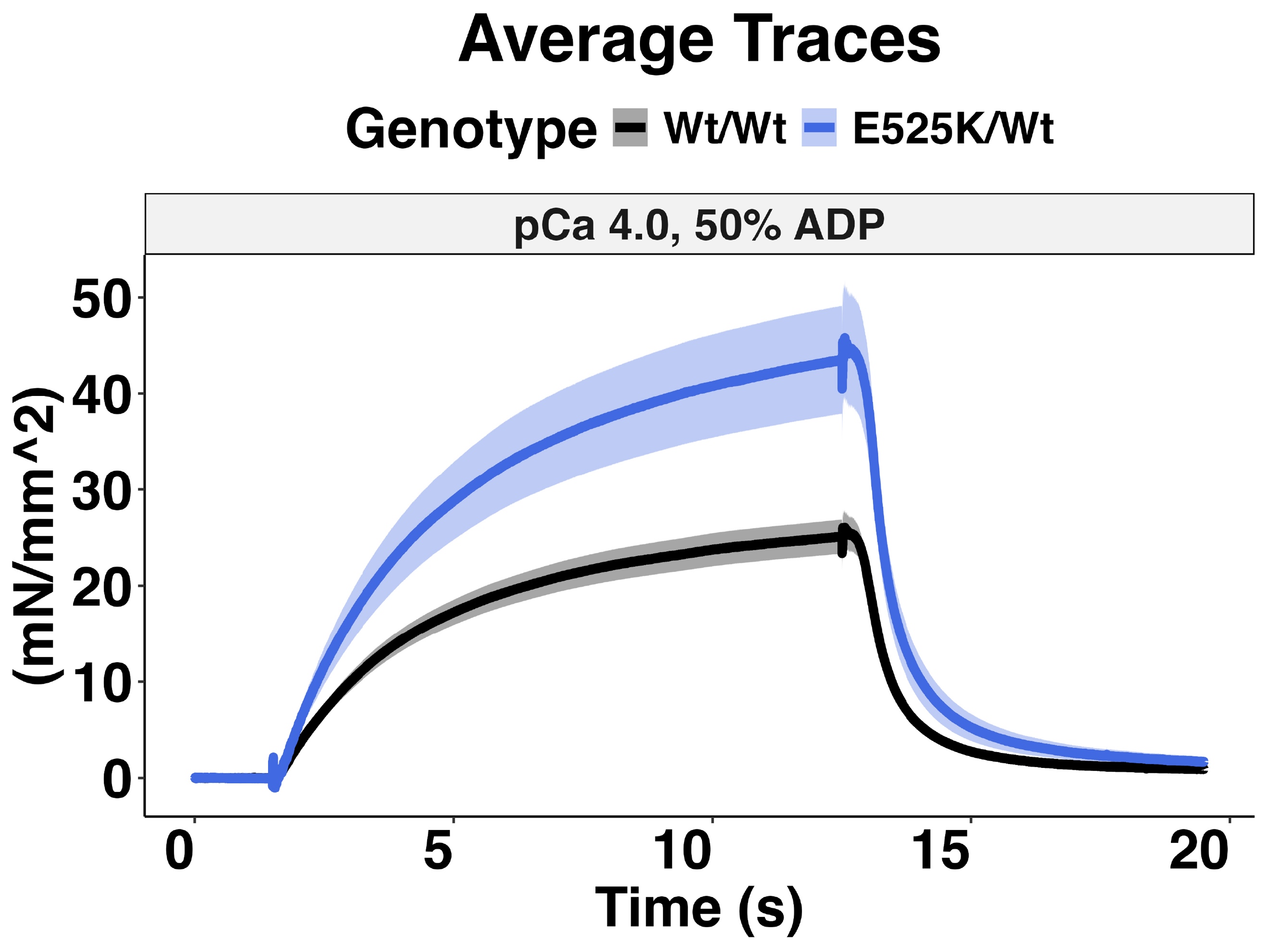

Supplemental Figure 8: Average unscaled myofibril trace in the presence of ADP

Average traces of normalized force over time (Mean +/- SEM) at maximal (pCa 4.0) stimulation.

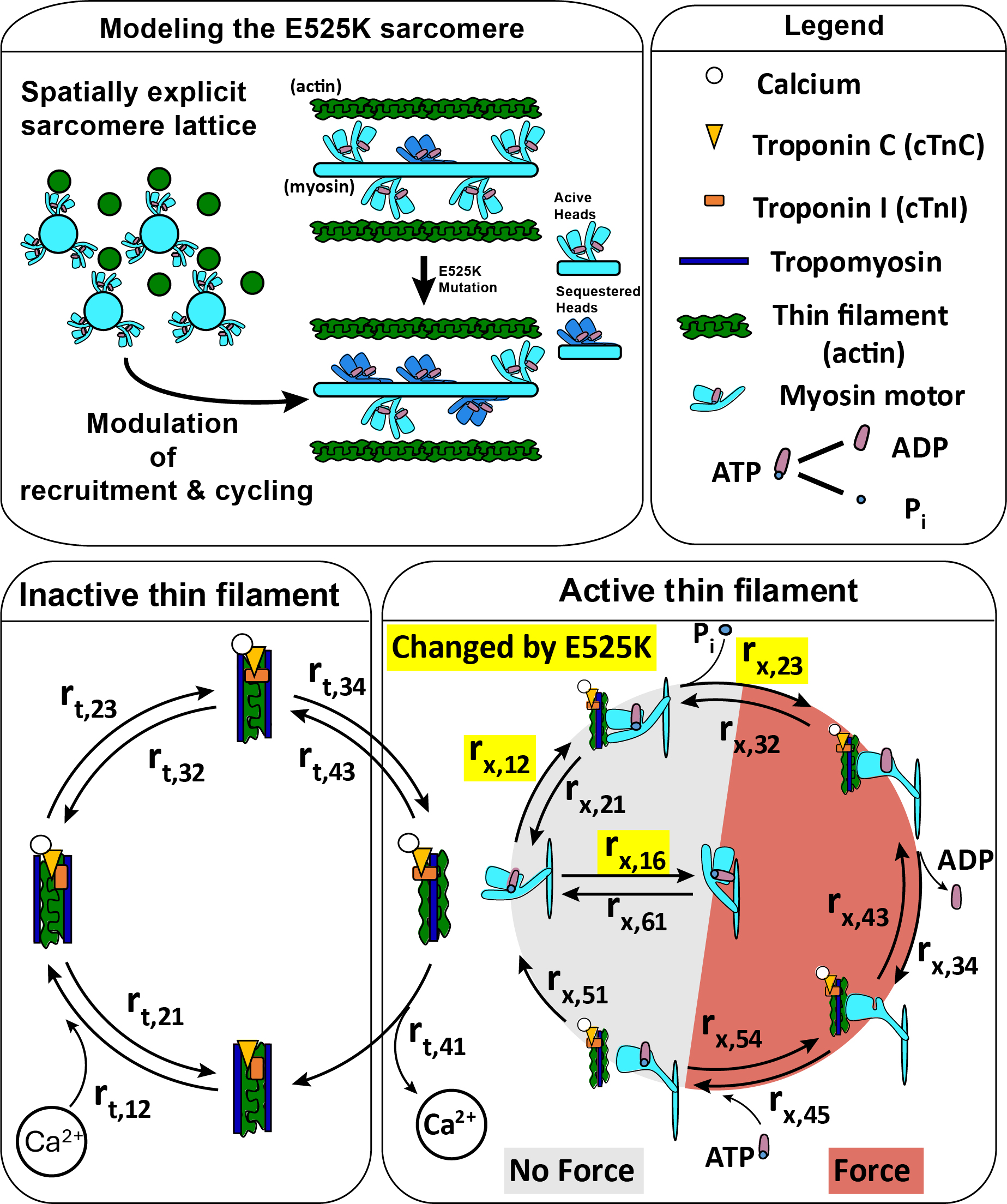

Supplemental Figure 9: Half sarcomere model

LEFT: Depiction of the sarcomere lattice, showing the thick (teal) and thin (green) filaments. Periodic boundary conditions are used so that each myosin and actin is fully surrounded by binding partners. In this model the thick and thin filaments are represented as networks of springs, with crossbridges modeled as paired torsional and linear springs. RIGHT: Rate transition diagram for the thick and thin filaments, showing the four thin-filament rates (r_t_) and six myosin motor rates (r_x_).

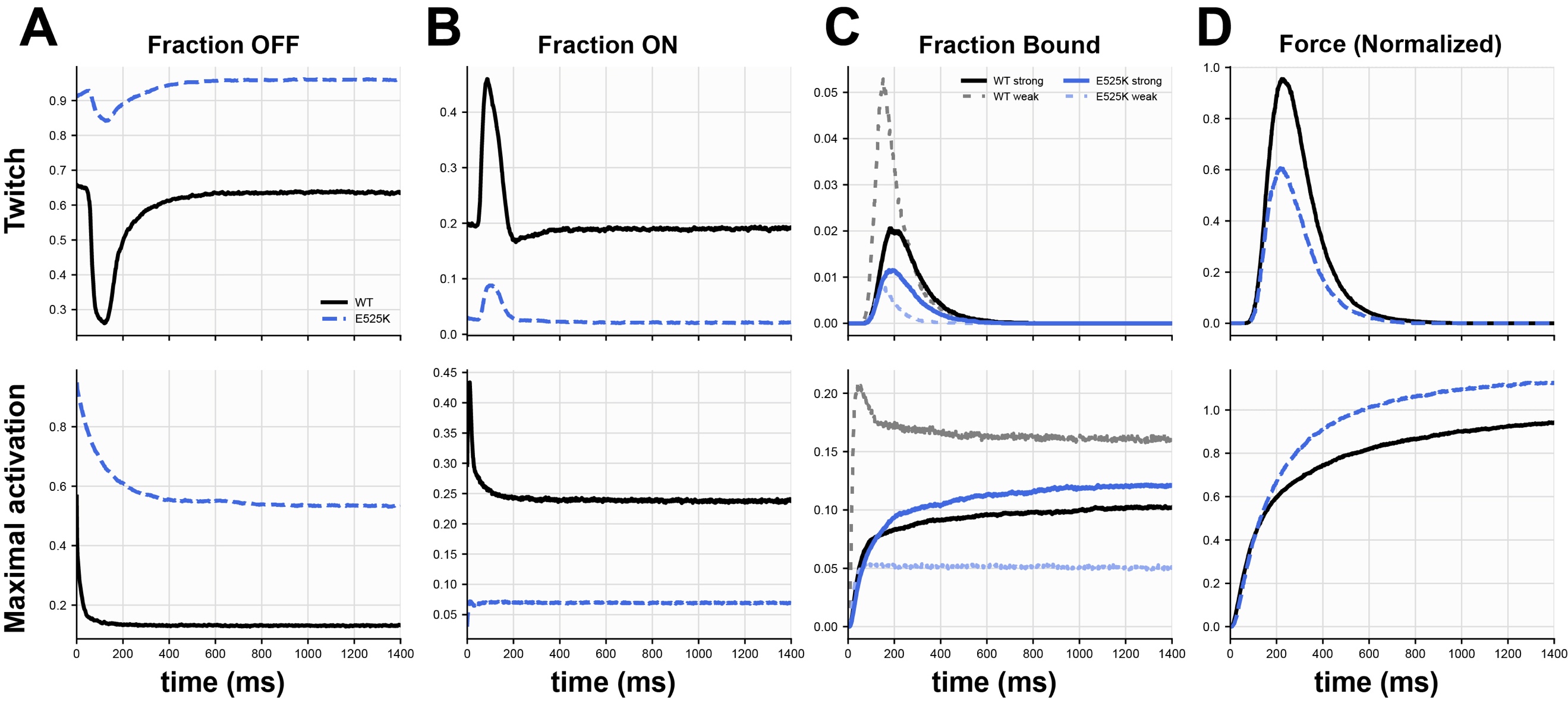

Supplemental Figure 10: Fraction of heads in the OFF, ON, weakly vs strongly bound, and the force, for twitch (upper row) and max isometric (lower row) simulations.

Solid lines indicate median, shaded error bars indicate inter-quartile range. The fraction of on heads is lower at every time step in the mutant than in the WT conditions. However, in max isometric simulations, in WT, the weakly bound portion rises rapidly and is takes about 200 ms to be reach steady state in fraction of strongly bound state, and the strongly bound portion never exceeds 10%. However, in the mutant, weakly bound cross-bridges rapidly convert to the strongly bound state, and the steady-state fraction is greater than that of the WT. In twitches, on the other hand, too few heads are recruited. The mutant has a greater fraction of strongly bound than weakly bound in the twitch, but not enough to overcome the head deficit.

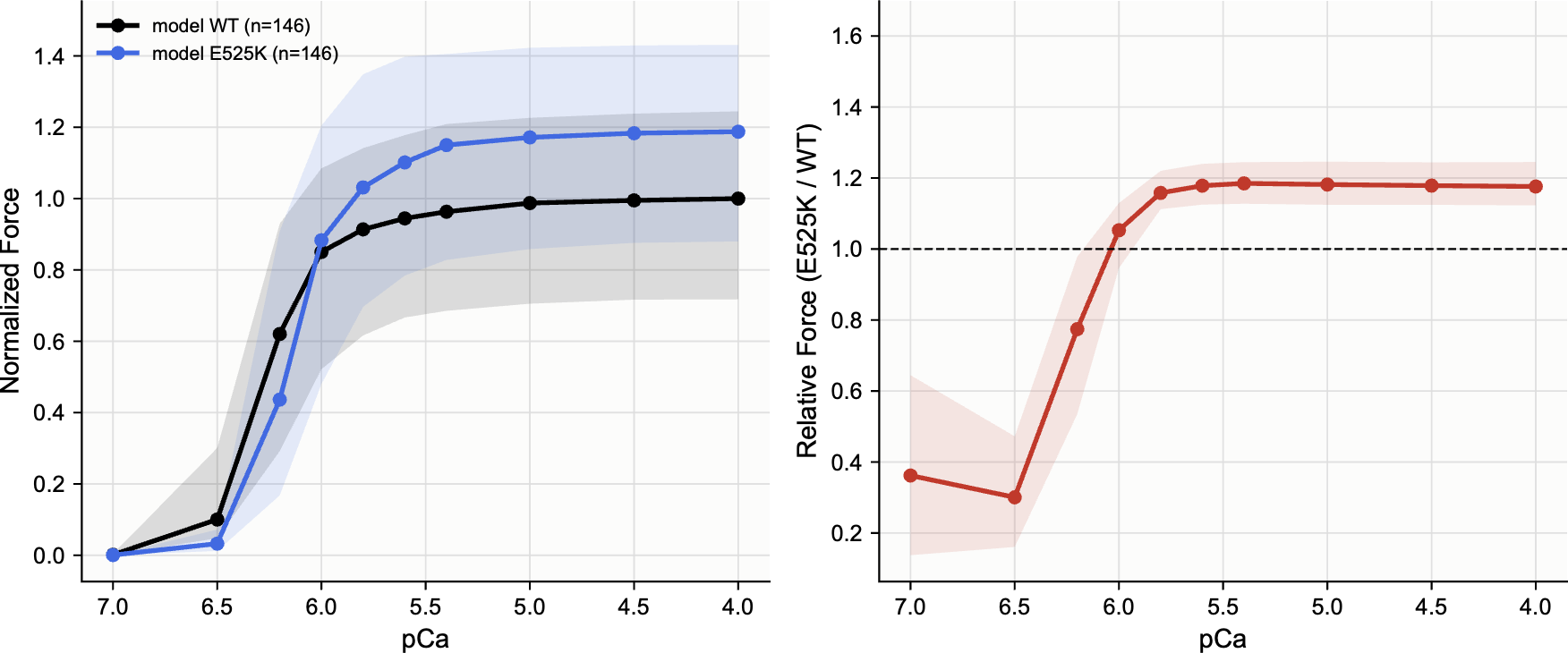

Supplemental Figure 11: Simulated force pCa curve

Left: simulated force-pCa curve, median, and interquartile range. The data comprise each baseline and the single perturbation that yielded the most E525K-like response. Right: relative E525K/WT force at each pCa. The mutation raises the steady-state force by a constant factor at each pCa, if the thin filament is fully activated.

### Supplemental Tables

**Supplemental Table 1:** Sampling ranges for the WT baseline parameter sweep

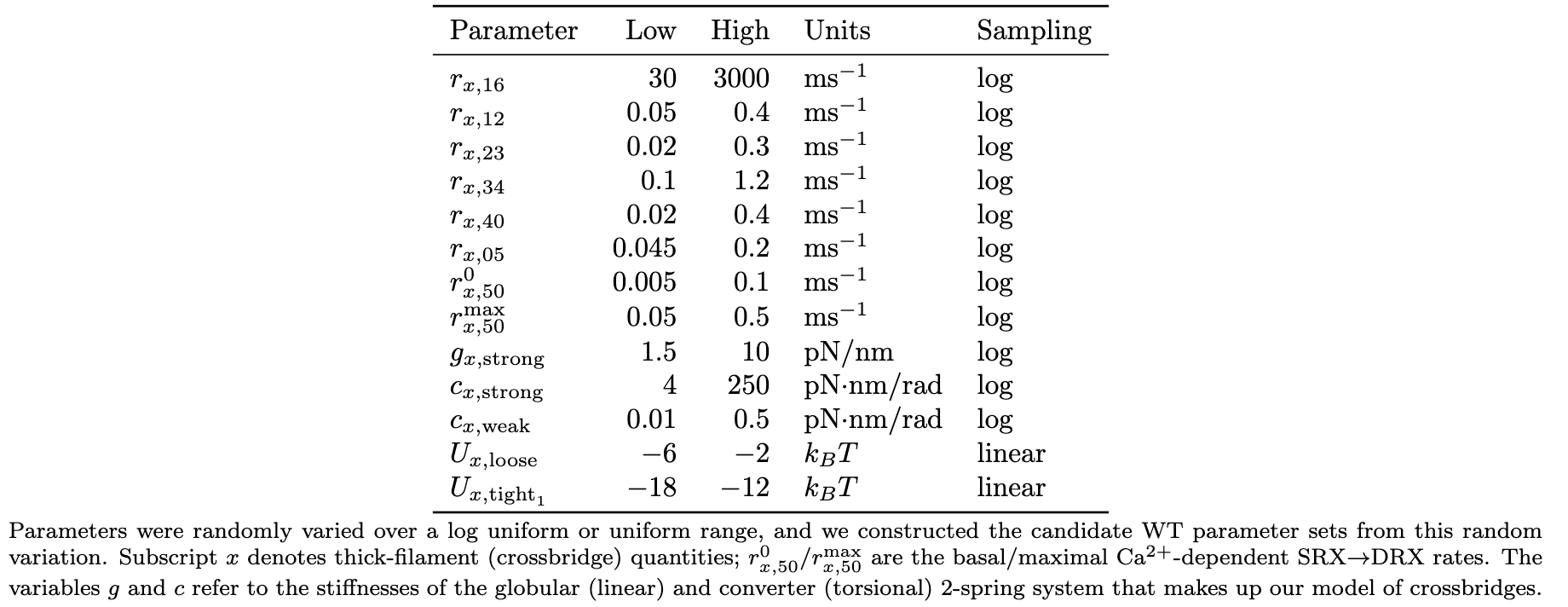

**Supplemental Table 2**: Simulated time constants

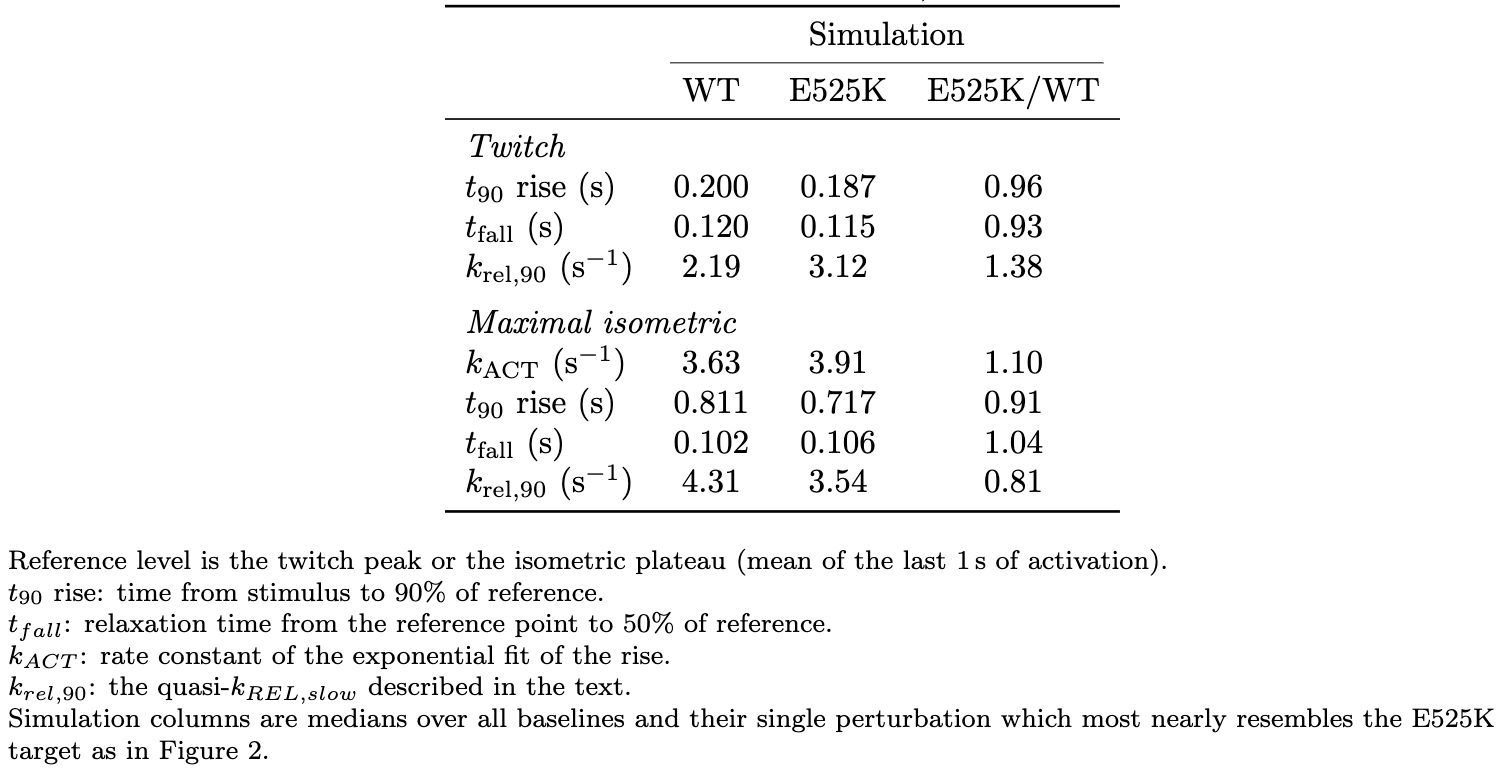

**Supplemental Table 3:** Crossbridge state occupancy and mean force per attached head, at the maximal-activation plateau and at the twitch peak
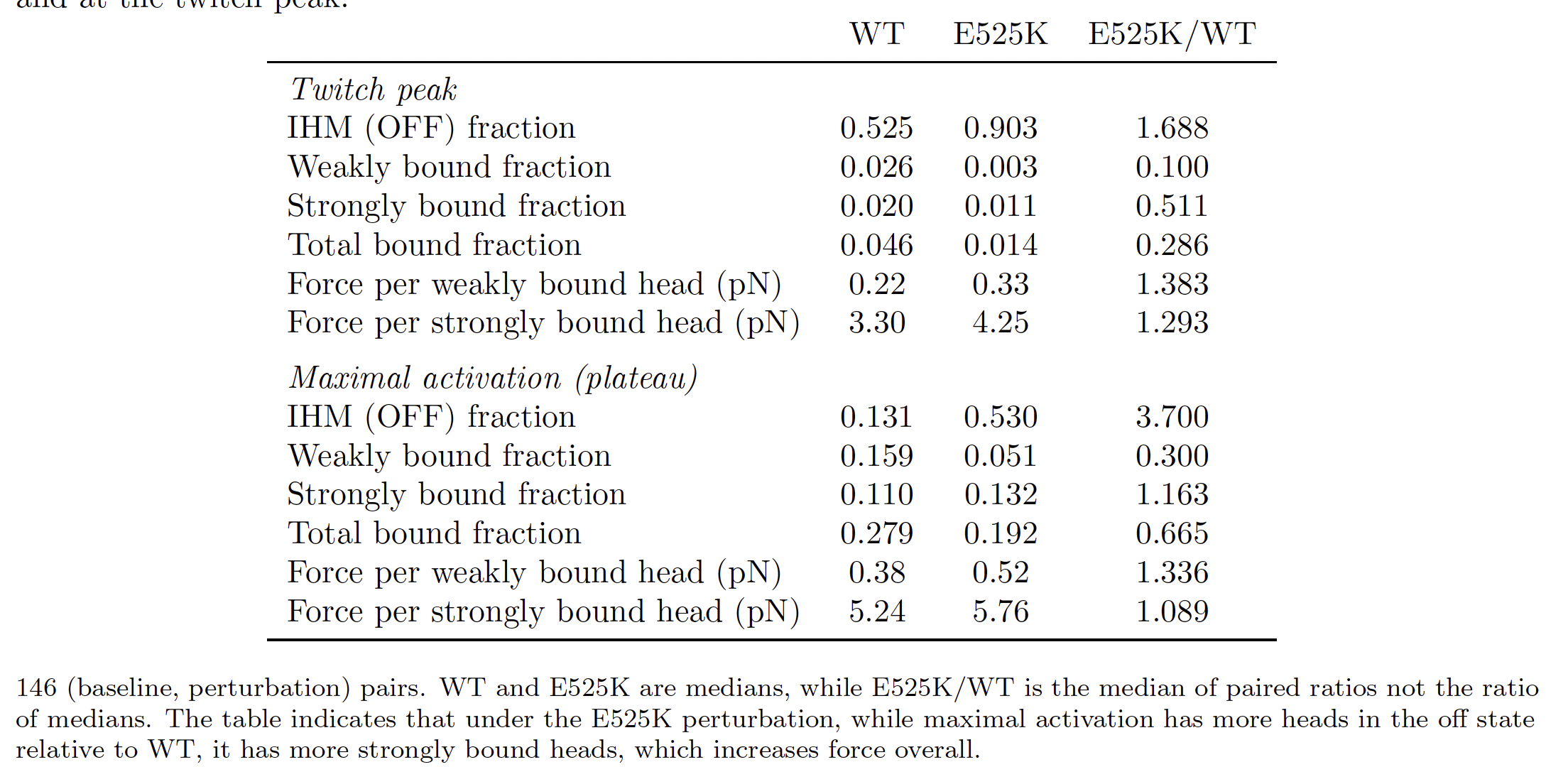

**Supplemental Table 4:** Impact of Pi release and OFF/IHM stabilization alone and together on k_ACT_

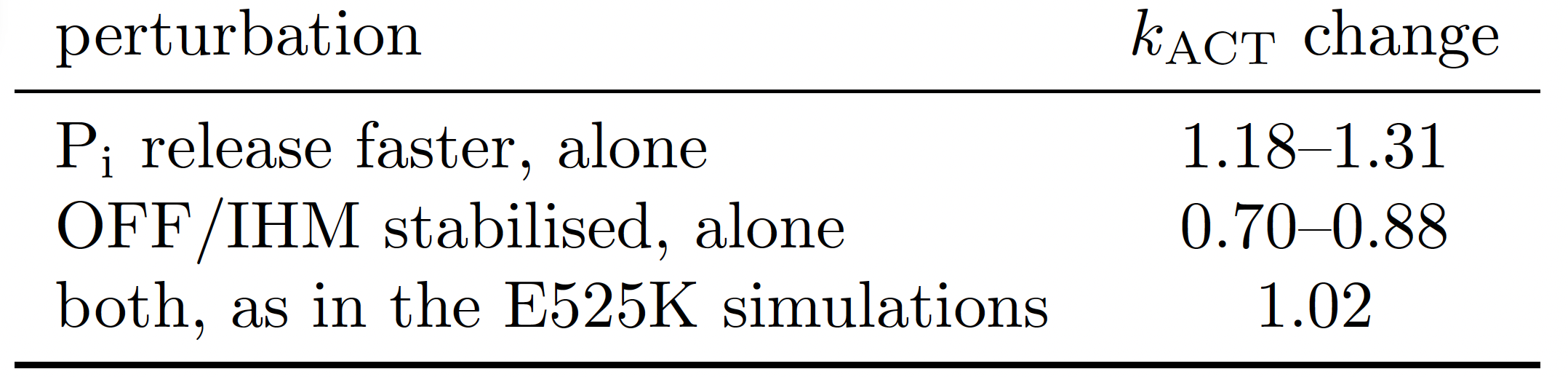
